# Two opposite voltage-dependent currents control the unusual early development pattern of embryonic Renshaw cell electrical activity

**DOI:** 10.1101/2020.06.18.158931

**Authors:** Juliette Boeri, Claude Meunier, Hervé Le Corronc, Pascal Branchereau, Yulia Timofeeva, François Xavier Lejeune, Christine Mouffle, Hervé Arulkandarajah, Jean Marie Mangin, Pascal Legendre, Antonny Czarnecki

**Author notes:** Corresponding authors (PL) (AC). These authors contributed equally to this work (Juliette Boeri, Claude Meunier, Hervé Le Corronc). These authors also contributed equally to this work (Pascal Legendre, Antonny Czarnecki).

## Abstract

Renshaw cells (V1^R^) are excitable as soon as they reach their final location next to the spinal motoneurons and are functionally heterogeneous. Using multiple experimental approaches, in combination with biophysical modeling and dynamical systems theory, we analyzed, for the first time, the mechanisms underlying the electrophysiological properties of V1R during early embryonic development of the spinal cord locomotor networks (E11.5-E16.5). We found that these interneurons are subdivided into several functional clusters from E11.5 and then display an unexpected transitory involution process during which they lose their ability to sustain tonic firing. We demonstrated that the essential factor controlling the diversity of the discharge pattern of embryonic V1^R^ is the ratio of a persistent sodium conductance to a delayed rectifier potassium conductance. Taken together, our results reveal how a simple mechanism, based on the synergy of two voltage-dependent conductances that are ubiquitous in neurons, can produce functional diversity in V1^R^ and control their early developmental trajectory.

## Introduction

The development of the central nervous system (CNS) follows complex steps, which depend on genetic and environmental factors and involve interactions between multiple elements of the neural tissue. Remarkably, emergent neurons begin to synchronize soon after the onset of synapse formation, generating long episodes of low frequency (<0.01 Hz) correlated spontaneous network activity (SNA) [1-8]. In the mouse embryonic spinal cord (SC), SNA is driven by an excitatory cholinergic-GABAergic loop between motoneurons (MNs) and interneurons (INs), GABA being depolarizing before embryonic day 16.5 (E16.5) [9]. SNA emerges around E12.5 [4, 6, 10-12], at a time when functional neuromuscular junctions are not yet established [13], and sensory and supraspinal inputs have not yet reached the spinal motor networks [14-17].

Several studies pointed out that SNA is an essential component in neuronal networks formation. [18-21]. In the SC, pharmacologically-induced disturbances of SNA between E12.5 and E14.5 induce defects in the formation of motor pools, in motor axon guidance to their target muscles and in the development of motor networks [4, 21-23]. During SNA episodes, long lasting giant depolarization potentials (GDPs) are evoked in the SC, mainly by the massive release of GABA onto MNs [12]. Immature Renshaw cells (V1^R^) are likely the first GABAergic partners of MNs in the mouse embryo [24, 25], and the massive release of GABA during SNA probably requires that many of them display repetitive action potential firing or plateau potential activity [25].

However, little is known about the firing pattern of embryonic V1^R^ and the maturation of their intrinsic properties. We recently found that V1^R^ exhibit heterogeneous excitability properties when SNA emerges in the SC [25] in contrast to adult Renshaw cells that constitute a functionally homogeneous population [26, 27]. Whether this early functional diversity really reflects distinct functional classes of V1^R^, how this diversity evolves during development, and what are the underlying biophysical mechanisms remain open questions. The present study addresses these issues using multiple approaches, including patch-clamp recordings, cluster analysis, biophysical modeling and dynamical systems theory. The firing patterns of V1^R^ and the mechanisms underlying their functional diversity are analyzed during a developmental period covering the initial phase of development of SC activity in the mouse embryo (E11.5-E14.5), when SNA is present, and during the critical period (E14.5-E16.5), when GABAergic neurotransmission gradually shifts from excitation to inhibition [28] and locomotor-like activity emerges [4, 10, 11].

We discover that the balance between the slowly inactivating subthreshold persistent sodium inward current (*I_Nap_*) [29] and the delayed rectifier potassium outward current (*I*_*Kdr*_), accounts for the heterogeneity of embryonic V1^R^ and the changes in firing pattern during development. The heterogeneity of V1^R^ at E12.5 arises from the existence of distinct functional groups. Surprisingly, and in opposition to the classically accepted development scheme [30-35], we show that the embryonic V1^R^ population loses its ability to support tonic firing from E13.5 to E15.5, exhibiting a transient functional involution during its development. Our experimental and theoretical results provide a global view of the developmental trajectories of embryonic V1^R^. They demonstrate that a simple mechanism, based on the synergy of only two major opposing voltage-dependent currents, accounts for functional diversity in these immature neurons.

## Results

### The delayed rectifier potassium current **I_Kdr_** is a key partner of the persistent sodium current **I_*Nap*_** in controlling embryonic V1^R^ firing patterns during development

We previously highlighted that V1^R^ are spontaneously active at E12.5. Their response to a 2 s suprathreshold depolarizing current steps revealed four main patterns, depending of the recorded interneuron [25]: i) single spiking (SS) V1^R^ that fires only 1-3 APs at the onset of the depolarizing pulse, ii) repetitive spiking (RS) V1^R^, iii) mixed events (ME) V1^R^ that shows an alternation of action potentials (APs) and plateau potentials or, iv) V1^R^ that displays a long-lasting sodium-dependent plateau potential (PP) (***Figure 1A1–A4***).

**Figure 1.**
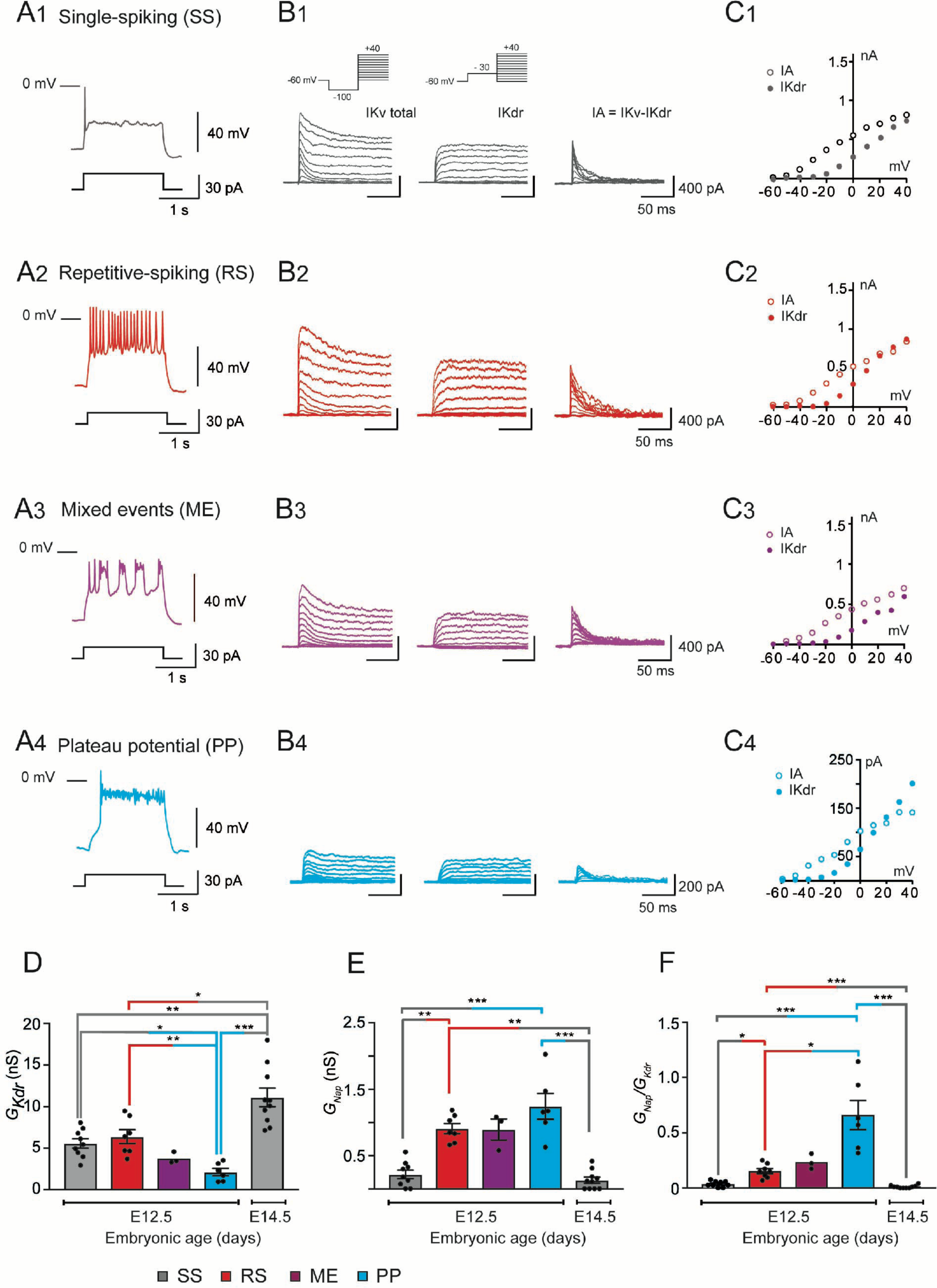
G_Kdr_ and G_Nap_ in embryonic V1^R^ at E12.5 and E14.5. (A) Representative traces of voltage responses showing single-spiking activity in E12.5 SS V1^R^ (A1), repetitive action potential firing in RS V1^R^ (A2), Mixed of plateau potential activity and repetitive action potential firing in ME V1^R^ (A3) and plateau potential activity in PP V1^R^ (A4). (B) Representative examples of the total outward K^+^ currents (IKV total) obtained from *V*_*H*_ = -100 mV (left traces), of *I*_*Kdr*_(*V*_*H*_ = -30 mV, middle traces) and of isolated *I*_*A*_ (left traces) recorded at E12.5 in SS V1^R^ (B1), RS V1^R^ (B2), ME V1^R^ (B3) and PP V1^R^ (B4). Voltage-dependent potassium currents were evoked in response to 10 mV membrane potential steps (200 ms) from -100 or from -30 mV to +40 mV (10 s interval between pulses). V1 ^R^ were voltage clamped at *V*_*H*_ = -60 mV. A prepulse of -40 mV (300 ms) was applied to activate both *I*_*A*_ and *I_Kdr_*.*I*_*Kdr*_ was isolated by applying a prepulse of 30 mV (300 ms) to inactivate *I*_*A*_ (B1 insert). *I*_*A*_ was isolated by subtracting step-by-step the currents obtained using a pre-pulse of 30 mV (*V*_*H*_ = - 30 mV) from the currents obtained using a pre-pulse of -40 mV (*V*_*H*_ = -100 mV). (C) Current-voltage relationship (*I* − *V* curves) of *I*_*Kdr*_ (filled circles) and of *I*_*A*_ (open circles) recorded in SS V1^R^ (C1), RS V1^R^ (C2), ME V1^R^ (C3) and PP V1^R^ (C4). *I* − *V* curves were obtained from currents shown in B1, B2, B3 and B4. Note that *I* − *V* curves are similar between SS V1^R^, RS V1^R^, ME V1^R^ and PP V1^R^. (D) Bar graph showing maximal *G*_*Kdr*_ value (Max *G*_*Kdr*_) in SS V1^R^ at E12.5 (n = 9; N = 9; gray bar) and at E14.5 (n = 10; N = 10 gray bar), and in RS V1^R^ (n = 7; N = 7; red bar), ME V1^R^ (n = 3; N = 3 purple bar) and PP V1^R^ at E12.5 (n = 7; N = 7 blue bar). *G*_*Kdr*_ was calculated from *I*_*Kdr*_ at *V*_*H*_ = + 20 mV, assuming a K^+^ equilibrium potential of -96 mV. There is no significant difference in *G*_*Kdr*_ between SS V1^R^ and RS V1^R^, while *G*_*Kdr*_ is significantly smaller in PP V1^R^ as compared to SS V1^R^ and RS V1^R^. *G*_*Kdr*_ was significantly higher in SS V1^R^ at E14.5 than in SS V1^R^, RS V1^R^ and PP V1^R^ at E12.5. (Kruskall-Wallis test *P* < 0.0001; SS V1^R^ versus RS V1^R^ at E12.5, *P* = 0.5864; SS V1^R^ versus PP V1^R^ at E12.5, *P* = 0.0243; RS V1^R^ versus PP V1^R^ at E12.5, *P* = 0.0086; E14.5 SS V1^R^ versus E12.5 SS V1^R^, *P* = 0.0048; E14.5 SS V1^R^ versus E12.5 RS V1^R^, *P* = 0.0384, E14.5 SS V1^R^ versus E12.5 PP V1^R^, *P* < 0.0001). The increase in *G*_*Kdr*_ between E12.5 and E14.5 is likely to be due to the increase in neuronal size (input capacitance; Figure 2A). Indeed, there was no significant difference (Mann Whitney test, *P* = 0.133) in *G*_*Kdr*_ density between SS V1^R^ at E12.5 (n = 9; N = 9 gray bar) and at E14.5 (n = 10; N = 10 gray bar). (E) Bar graph showing the maximal *G*_*Nap*_ value (Max *G*_*Nap*_) in SS V1^R^ at E12.5 (n = 9; N = 9 gray bar) and E14.5 (n = 10; N = 10 gray bar), and in RS V1^R^ (n = 8; N = 8 red bar), ME V1^R^ (n = 3; N = 3 purple bar) and PP V1^R^ (n = 6; N = 6 blue bar) at E12.5. Max *G*_*Nap*_ was calculated from maximal *I*_Nap_ value measured on current evoked by assuming a Na^+^ equilibrium potential of +60 mV. There was no difference in *G*_*Nap*_ between RS V1^R^ and PP V1^R^. On the contrary, *G*_*Nap*_ measured in SS V1^R^ at E12.5 or at E14.5 was significantly smaller as compared to *G*_*Nap*_ measured at E12.5 in RS V1^R^ or in PP V1^R^. *G*_*Nap*_ measured at E12.5 and E14.5 in SS V1^R^ were not significantly different (Kruskall-Wallis test *P* < 0.0001; E12.5 SS V1^R^ versus E12.5 RS V1^R^, *P* = 0.0034; E12.5 SS V1^R^ versus E12.5 PP V1^R^, *P* = 0.0006; E12.5 RS V1^R^ versus E12.5 PP V1^R^, *P* = 0.5494; E14.5 SS V1^R^ versus E12.5 SS V1^R^, *P* = 0.5896; E14.5 SS V1^R^ versus E12.5 RS V1^R^, *P* = 0.0005; E14.5 SS V1^R^ versus E12.5 PP V1^R^, *P* < 0.0001). (F) Histograms showing the *G*_*Nap*_/*G*_*Kdr*_ ratio in SS V1^R^ at E12.5 (n = 9; gray bar) and E14.5 (n = 10; green bar) and in RS V1^R^ (n = 8; red bar), ME V1^R^ (n = 3; purple bar) and PP V1^R^ (n = 6; blue bar) at E12.5. Note that the *G*_*Nap*_/*G*_*Kdr*_ ratio differs significantly between SS V1^R^, RS V1^R^ and PP V1^R^ at E12.5, while it is not different between SS V1^R^ recorded at E12.5 and at E14.5 (Kruskall-Wallis test *P* < 0.0001; SS V1^R^ versus RS V1^R^ at E12.5, *P* = 0.0367; SS V1^R^ versus PP V1^R^ at E12.5, *P* < 0.0001; RS V1^R^ versus PP V1^R^ at E12.5, *P* = 0.0159; E14.5 SS V1^R^ versus E12.5 SS V1^R^, *P* = 0.2319; E14.5 SS V1^R^ versus E12.5 RS V1^R^, *P* = 0.0017; E14.5 SS V1^R^ versus E12.5 PP V1^R^ *P* < 0.0001). Data shown in A and B were used to calculate *G*_*Nap*_/*G*_*Kdr*_ ratio shown in C. (\**P* < 0.05, ** *P* < 0.01, *** *P* < 0.001).

We also uncovered a relationship between *I*_*Nap*_ and the ability of embryonic V1^R^ to sustain repetitive firing [25]. However, the heterogeneous firing patterns of V1^R^ observed at E12.5 could not be fully explained by variations in *I*_*Nap*_[25], suggesting the involvement of other voltage-gated channels in the control of the firing pattern of V1^R^, in particular potassium channels, known to control firing and AP repolarization. Our voltage clamp protocol, performed in the presence of TTX (1 μM), did not disclose any inward rectifying current (hyperpolarizing voltage steps to -100 mV from *V*_*H*_ = -20 mV, data not shown), but revealed two voltage-dependent outward potassium currents, a delayed rectifier current (*I*_Kdr_) and a transient potassium current (*I*_*A*_) in all embryonic V1^R^, whatever the firing pattern (***Figure 1B1–B4***). These currents are known to control AP duration (*I*_*Kdr*_) or firing rate (*I_A_*), respectively [36]. The activation threshold of *I*_*Kdr*_ lied between -30 mV and -20 mV and the threshold of *I*_*A*_ between -60 mV and -50 mV, (n = 27; N = 27 embryos) (***Figure 1C1–C4***). Removing external calcium had no effect on potassium current I/V curves (data not shown), suggesting that calcium-dependent potassium currents are not yet present at E12.5.

It was unlikely that the heterogeneity of V1^R^ firing patterns resulted from variations in the intensity of *I_A_*. Indeed, its voltage-dependent inactivation (time constant: 23.3 ± 2.6 ms, n = 8; N = 8), which occurs during the depolarizing phase of an AP, makes it ineffective to control AP or plateau potential durations. This was confirmed by our theoretical analysis (***see Figure 7––figure supplement 1***). We thus focused our study on *I*_*Kdr*_. At E12.5, PP V1^R^ had a significantly lower *G*_*Kdr*_(2.12 ± 0.44 nS, n = 6; N = 6) than SS V1^R^ (5.57 ± 0.56 nS, n = 9; N = 9) and RS V1^R^ (6.39 ± 0.83 nS, n = 7; N = 7) (***Figure 1D***). However, there was no significant difference in *G*_*Nap*_ between SS V1^R^ and RS V1^R^ at E12.5 (***Figure 1D***), which indicated that variations in *G*_*Nap*_ alone could not explain all the firing patterns observed at E12.5. Similarly, there was no significant difference in *G*_*Kdr*_ between RS V1^R^ (0.91 ± 0.21nS, n = 8; N = 8) and PP V1^R^ (1.24 ± 0.19 nS, n = 6; N = 6) at E12.5 (***Figure 1E***), indicating that variations in *G*_*Kdr*_ alone could not explain all the firing patterns of V1^R^ at E12.5 [25]. In contrast *I*_*A*_ measured in SS V1^R^ at E12.5 (0.21 ± 0.20 nS, n = 9; N = 9) were significantly lower compared to *G*_*Nap*_ measured in RS V1^R^ and in PP V1^R^ at E12.5 (***Figure 1E***).

Mature neurons often display multiple stable firing patterns [37-39]. This usually depends on the combination of several outward and inward voltage- or calcium-dependent conductances and on their spatial localization [37-39]. In contrast, immature V1^R^ have a limited repertoire of voltage-dependent currents (*I*_*Nat*_ and *I*_*Nap*_, *I*_*Kdr*_ and *I*_*A*_) at E12.5, and we did not find any evidence of voltage-dependent calcium currents at this age [25]. Blocking *I*_*Nap*_ prevented plateau potential activity, PP-V1^R^ becoming unexcitable, and turned repetitive spiking V1^R^ into single spiking V1^R^ [25]. Therefore, we hypothesized that the different firing patterns of V1^R^ observed at E12.5 were related to the *G*_*Nap*_/*G*_*Kdr*_ ratio only, with variations in the intensity of *I*_*A*_ being unlikely to account for the heterogeneity of firing pattern. We found that this ratio was significantly lower for SS V1^R^ recorded at E12.5 (*G*_*Nap*_/*G*_*Kdr*_= 0.043 ± 0.015, n = 9) compared to RS V1^R^ (0.154 ± 0.022, n = 8) and PP V1^R^ (0.66 ± 0.132, n = 6) (***Figure 1F***). We also found that the *G*_*Nap*_/ *G*_*Kdr*_ ratio was significantly lower for RS V1^R^ compared to PP V1^R^ (***Figure 1F***).

Altogether, these results strongly suggest that, although the presence of *I*_*Nap*_ is required for embryonic V1^R^ to fire repetitively or to generate plateau potentials [25], the heterogeneity of the firing pattern observed between E12.5 is not determined by *I*_*Nap*_ *per se* but likely by the balance between *I*_*Nap*_ and *I*_*Kdr*_.

### Manipulating the balance between G_Nap_ and G_Kdr_ changes embryonic V1^R^ firing patterns

We previously showed that blocking *I*_*Nap*_ with riluzole converted PP V1^R^ or RS V1^R^ into SS V1^R^ [25]. To confirm further that the balance between *G*_*Nap*_ and *G*_*Kdr*_ was the key factor in the heterogeneity of V1^R^ firing patterns, we assessed to what extent a given E12.5 SS V1^R^ cell could change its firing pattern when *I*_*Kdr*_ was gradually blocked by 4-aminopiridine (4-AP). We found that *I*_*Kdr*_ could be blocked by micromolar concentrations of 4-AP without affecting *I_A_* (***Figure 2––figure supplement 1***). 4-AP, applied at concentrations ranging from 0.3 μM to 300 μM, specifically inhibited *I*_*Kdr*_ with an IC_50_ of 2.9 μM (***Figure 2––figure supplement 1C1***).

**Figure 2.**
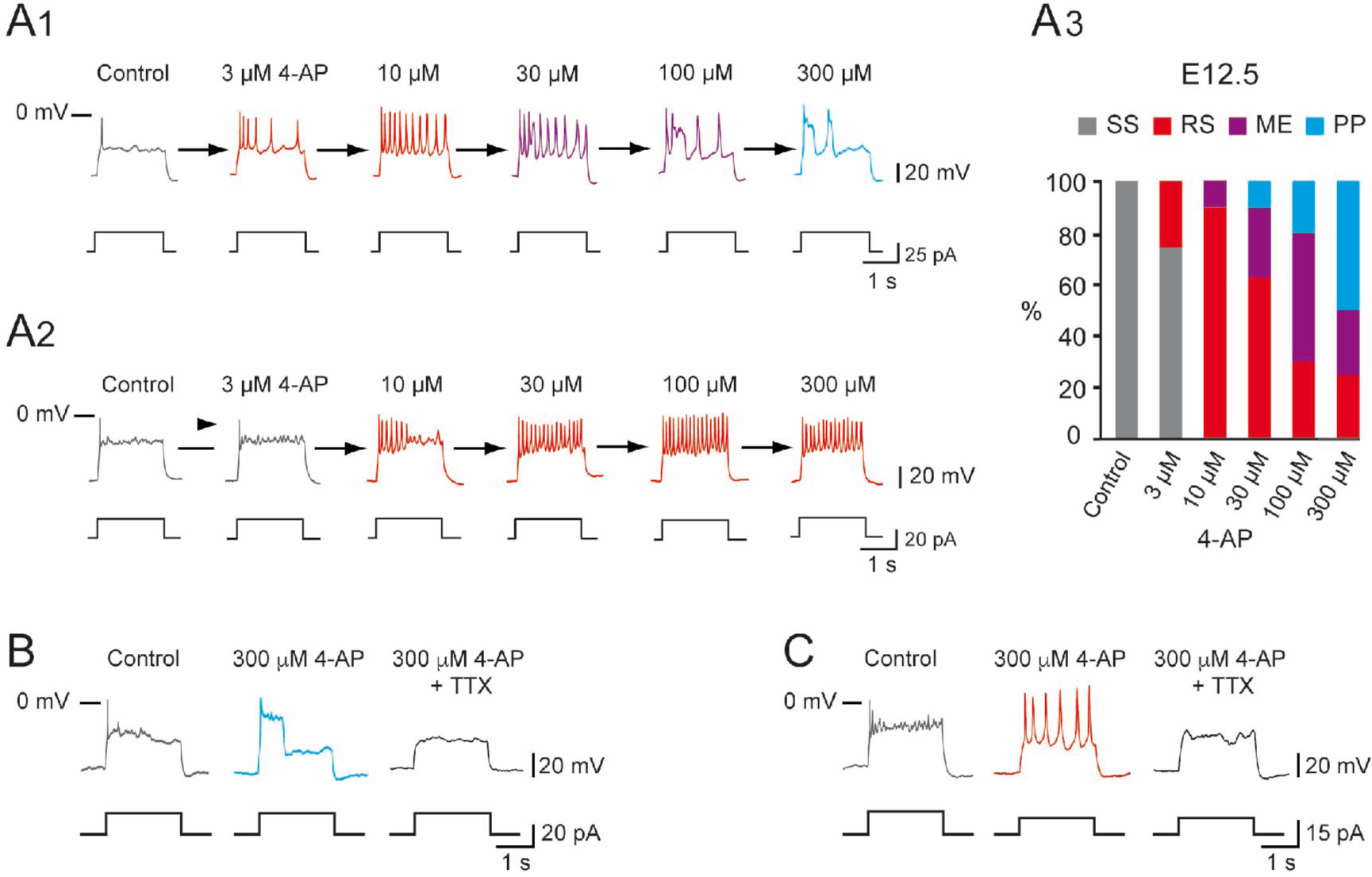
Increasing 4-AP concentration changed the firing pattern of single spiking embryonic V1^R^ recorded at E12.5. The firing pattern of embryonic V1^R^ was evoked by 2 s suprathreshold depolarizing current steps. (A) Representative traces showing examples of the effect of increasing concentration of 4-AP (from 3 to 300 μM) on the firing pattern of a SS V1^R^ recorded at E12.5. Note that in A1, increasing 4-AP concentration converted single spiking (gray trace) to repetitive spiking (red trace), repetitive spiking to a mixed event pattern (purple trace) and mixed events to plateau potential (blue trace). (A2) Example of SS V1^R^ in which increasing 4-AP concentration converted single spiking to repetitive spiking only. (A3) Bar plots showing the change in the firing pattern of SS V1^R^ according to 4-AP concentrations (control n = 10; N = 10, 3 μM 4-AP n = 8; N = 8, 10 μM 4-AP n = 10; N = 10, 30 μM 4-AP n = 10; N = 10, 100 μM 4-AP n = 10; N = 10, 300 μM 4-AP n = 8; N = 8). (B) Representative traces showing the effect of 0.5 μM TTX on a plateau potential evoked in a SS V1^R^ in the presence of 300 μM 4-AP. (C) Representative traces showing the effect of 0.5 μM TTX on repetitive AP firing evoked in a SS V1^R^ in the presence of 300 μM 4-AP. In both cases, the application of TTX fully blocked the responses evoked in the presence of 4-AP, indicating that they were underlain by the activation of voltage-gated Na^+^ channels.

We then determined to what extent increasing the concentration of 4-AP modified the firing pattern of V1^R^ at E12.5. Applying 4-AP at concentrations ranging from 3 μM to 300 μM changed the firing pattern of SS V1^R^ (n = 10; N = 10) in a concentration-dependent manner (***Figure 2A1–A3***). In 50% of the recorded V1^R^, increasing 4-AP concentrations successfully transformed SS V1^R^ into PP V1^R^ with the following sequence: SS → RS → ME → PP (***Figure 2A1***). In a second group of embryonic V1^R^ (25%), 4-AP application only evoked mixed activity, with the same sequence as aforementioned (SS → RS → ME) (data not shown). In the remaining SS V1^R^ (25%), increasing 4-AP concentration only led to sustained AP firing (***Figure 2A2***). Application of 300 μM 4-AP on RS V1^R^ at E12.5 evoked mixed events or plateau potentials (***Figure 2––figure supplement 2***). Plateau potentials and repetitive spiking evoked in the presence of 300 μM 4-AP were fully blocked by 0.5-1 μM TTX, indicating that they were generated by voltage-gated Na^+^ channels (***Figure 2B,C*** and ***Figure 2–figure supplement 2***). It should be noted that the application of 300 μM of 4-AP induced a significant 30.5 ± 12.4 % increase (*P* = 0.0137; Wilcoxon test) of the input resistance (1.11 ± 0.08 GΩ versus 1.41 ± 0.12 GΩ; n = 11; N = 11).

These results show that, in addition to *I*_*Nap*_, *I*_*Kdr*_ is also a major determinant of the firing pattern of embryonic V1^R^. The above suggests that the firing patterns depend on a synergy between *I*_*Nap*_ and *I*_*Kdr*_ and that the different patterns can be ordered along the following sequence SS → RS → ME → PP when the ratio *G*_*Nap*_/ *G*_*Kdr*_ is increased.

### The heterogeneity of the **V1^R^** firing patterns decreases during embryonic development

It was initially unclear whether these different firing patterns corresponded to well separated classes within the E12.5 V1^R^ population or not. To address this question, we performed a hierarchical cluster analysis on 163 embryonic V1^R^, based on three quantitative parameters describing the firing pattern elicited by the depolarizing pulse: the mean duration of evoked APs or plateau potentials measured at half-amplitude (mean ½Ad), the variability of the event duration during repetitive firing (coefficient of variation of ½Ad: CV ½Ad) and the total duration of all events, expressed in percentage of the pulse duration (depolarizing duration ratio: ddr) (***Figure 3A inserts***). In view of the large dispersion of mean ½Ad and ddr values, cluster analysis was performed using the (decimal) logarithm of these two quantities [40]. The analysis of the distribution of log mean ½Ad, CV ½Ad and log ddr revealed multimodal histograms that could be fitted with several Gaussians (***Figure 3–– figure supplement 1A1–C1*)**. Cluster analysis based on these three parameters showed that the most likely number of clusters was 5 (***Figure 3A,B***), as determined by the silhouette width measurement (***Figure 3B***). Two clearly separated embryonic V1^R^ groups with CV ½Ad = 0 stood out, as shown in the 3D plot in ***Figure 5C***. The cluster with the largest ½Ad (mean ½Ad = 833.5 ± 89.99 ms) and the largest ddr (0.441 ± 0.044) contained all PP V1^R^ (n = 35; N = 29) (***Figure 3C-D*** and ***Figure 3–figure supplement 1A2,C2***). Similarly, the cluster with the shortest ½Ad (9.73 ± 0.66 ms) and the lowest ddr (0.0051 ± 0.0004) contained all SS V1^R^ (n = 46; N = 37) (***Figure 3C-D*** and ***Figure 3–figure supplement 1A2,C2***).

**Figure 3.**
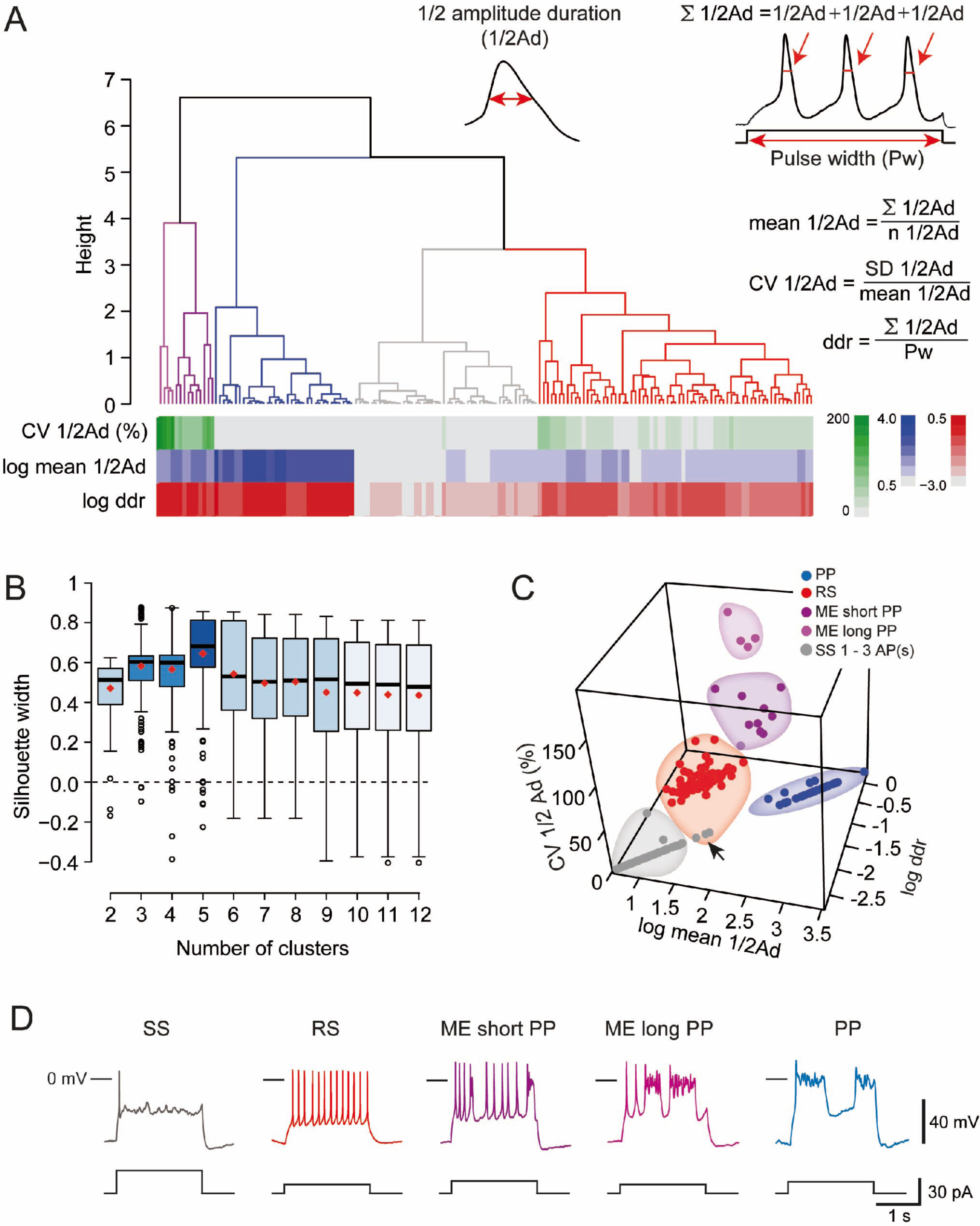
Cluster analysis of V1^R^ firing pattern at E12.5. (A, inserts) Cluster analysis of embryonic V1^R^ firing pattern was performed using three parameters that describe the firing pattern during a 2 s suprathreshold depolarizing pulses: the mean of the half-amplitude event duration (mean ½Ad), the coefficient of variation of ½ Ad (CV ½Ad) allowing to quantify the AP variation within a train (CV was set to 0 when the number of spikes evoked by a depolarizing pulse was ≤ 3) and the duration ratio ddr = Σ½ Ad/Pw, obtained by dividing the sum of ½ Ad by the pulse duration Pw, that indicates the total time spent in the depolarized state. For example, ddr = 1 when a plateau potential lasts as long as the depolarizing pulse. Conversely, its value is low when the depolarizing pulse evokes a single AP only. (A) Dendrogram for complete linkage hierarchical clustering of 164 embryonic V1^R^ (N = 140) according to the values of log mean ½Ad, of CV ½Ad and of log ddr. The colored matrix below the dendrogram shows the variations of these three parameters for all the cells in the clusters (colored trees) extracted from the dendrogram. (B) The number of clusters was determined by analyzing the distribution of silhouette width values (see Material and Methods). The boxplots show the distribution of silhouette width values when the number of clusters k varies from 2 to 12. The mean silhouette width values (red diamond shaped points) attained their maximum when the estimated cluster number was 5. (C) 3D plot showing cluster distribution of embryonic V1^R^ according to log mean ½Ad, CV ½Ad and log ddr. Each cluster corresponds to a particular firing pattern as illustrated in D. V1^R^ that cannot sustain repetitive firing of APs (1 to 3 AP/pulse only, gray, Single spiking, SS), V1^R^ that can fire tonically (red, Repetitive spiking, RS), V1^R^ with a firing pattern characterized by a mix of APs and relatively short plateau potentials (dark purple, Mixed event short PP, ME short PP), V1^R^ with a firing pattern characterized by a mix of APs and relatively long plateau potentials (light purple, Mixed event long PP, ME long PP) and V1^R^ with evoked plateau potentials only (blue, Plateau potential, PP). The arrow in C indicates 3 misclassified V1^R^ that could not sustain repetitive firing although they were assigned to the cluster of repetitively firing V1^R^ (see text).

The three other clusters corresponded to V1^R^ with nonzero values of CV ½Ad (***Figure 3C***). A first cluster regrouping all RS V1^R^ (n = 69; N = 61) was characterized by smaller values of ½Ad (23.91 ± 1.43 ms), CV ½Ad (27.36 ± 1.64%) and ddr (0.11 ± 0.01) (***Figure 3C-D*** and ***Figure 3–– figure supplement 1A2,C2***). The last two clusters corresponded to ME V1^R^ (***Figure 3C,D***). The smaller cluster, characterized by a larger CV ½Ad (170.9 ± 8.9%; n= 4; N = 4), displayed a mix of APs and short plateau potentials, while the second cluster, with smaller CV ½Ad (87.61 ± 7.37%; n = 9; N = 9), displayed a mix of APs and long-lasting plateau potentials (***Figure 3D*** and ***Figure 3–figure supplement 1B2***). Their ½Ad and ddr values were not significantly different (***Figure 3–figure supplement 1A2,C2***).

It must be noted that three embryonic V1^R^ (1.8%) were apparently misclassified since they were aggregated within the RS cluster although having zero CV ½Ad (***Figure 3C; arrows***). Examination of their firing pattern revealed that this was because they generated only two APs, although their ddr (0.16 to 0.2) and ½ Ad values (31.6 to 40.3 ms) were well in the range corresponding of the RS cluster.

These different firing patterns of V1^R^ might reflect different states of neuronal development [31, 41-43]. Single spiking and/or plateau potentials are generally believed to be the most immature forms of firing pattern, repetitive spiking constituting the most mature form [19, 44]. If it were so, the firing patterns of embryonic V1^R^ would evolve during embryonic development from single spiking or plateau potential to repetitive spiking, this latter firing pattern becoming the only one in neonates [26] and at early postnatal stages [27]. However, RS neurons already represent 41% of V1^R^ at E12.5. We therefore analyzed the development of firing patterns from E11.5, when V1^R^ terminate their migration and reach their final position [45], to E16.5. This developmental period covers a first phase of development (E11.5-E14.5), where lumbar spinal networks exhibit SNA, and a second phase (E14.5-E16.5), where locomotor-like activity emerges [4, 11, 46, 47]. We first analyzed changes in the intrinsic properties (input capacitance *C*_*in*_, input resistance *R*_*in*_ = 1/*G*_*in*_ and spike voltage threshold) of V1^R^. *C*_*in*_ did not change significantly from E11.5 to E13.5 (***Figure 4A1***), remaining of the order of 12 pF, in agreement with our previous work [25]. However, it increased significantly at the transition between the two developmental periods (E13.5-E15.5) to reach about 23.5 pF at E15.5 (***Figure 4A1***). A similar developmental pattern was observed for *R*_*in*_, which remained stable during the first phase from E11.5 to E14.5 (*R*_*in*_≈ 1-1.2 GΩ) but decreased significantly after E14.5 to reach about 0.7 GΩ at E15.5 (***Figure 4A2***).

**Figure 4.**
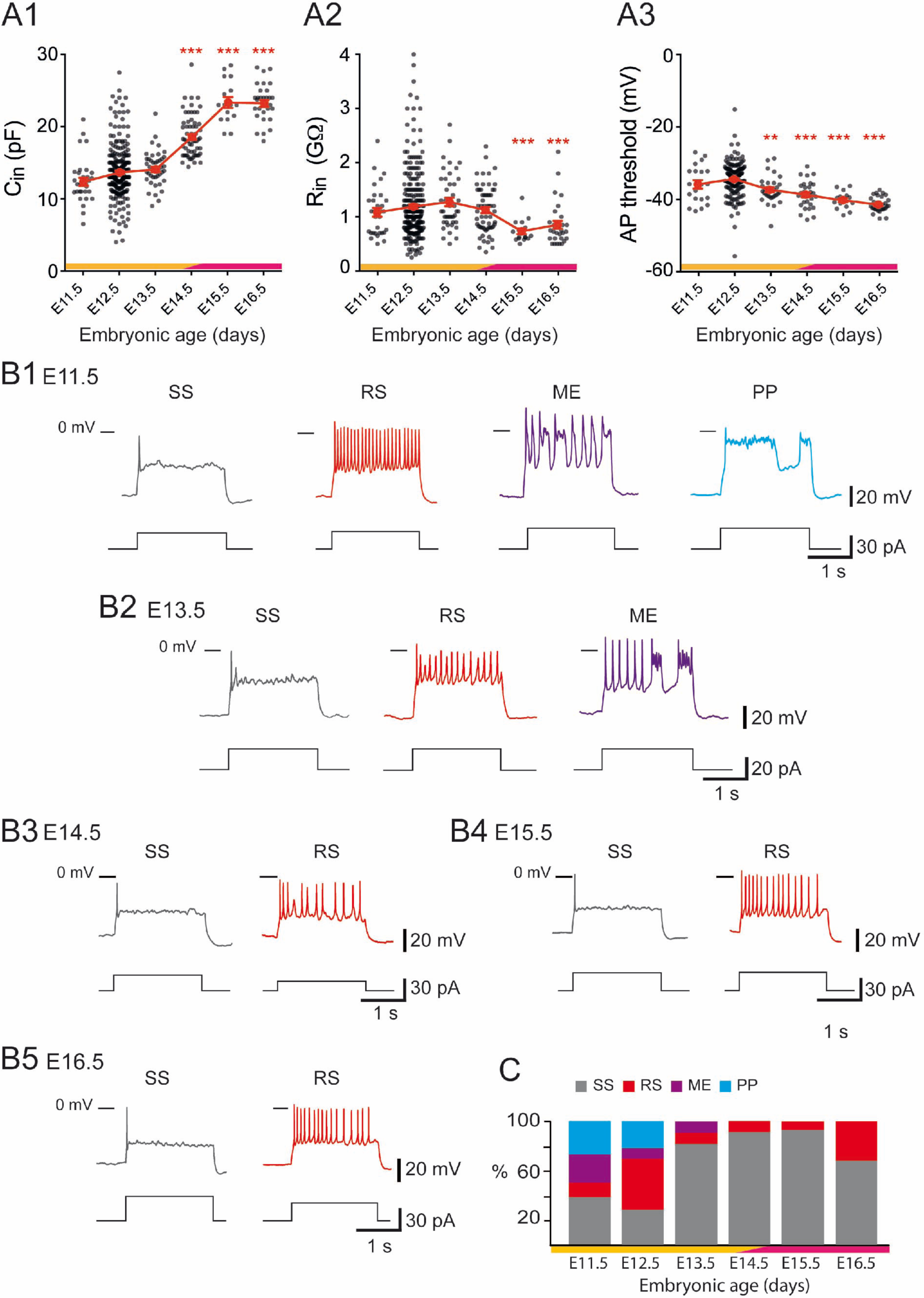
Developmental changes of embryonic V1^R^ firing patterns from E11.5 to E16.5. (A1) Graph showing how the input capacitance *C*_*in*_ of V1^R^ changes with embryonic age. *C*_*in*_ significantly increased between E12.5 or E13.5 and E14.5 (Kruskall-Wallis test *P* < 0.0001; E12.5 versus E11.5 *P* = 0.258, E12.5 versus E13.5 *P* = 0.904, E12.5 versus E14.5 *P* < 0.0001, E12.5 versus E15.5 *P* < 0.0001, E12.5 versus E16.5 *P* < 0.0001, E13.5 versus E14.5 *P* < 0.0001, E13.5 versus E15.5 *P* < 0.0001, E13.5 versus E16.5 *P* < 0.0001; E11.5 n = 31; N = 27, E12.5 n = 267; N = 152, E13.5 n = 43; N = 40, E14.5 n = 61; N = 49, E15.5 n = 16; N = 4, E16.5 n = 30; N = 9). (A2) Graph showing how the input resistance *R*_*in*_ of V1^R^ changes with embryonic age. *R*_*in*_ significantly decreased between E12.5 or E14.5 and E15.5 (Kruskall-Wallis test *P* < 0.0001; E12.5 versus E11.5 *P* > 0.999, E12.5 versus E13.5 *P* = 0.724, E12.5 versus E14.5 *P* > 0.999, E12.5 versus E15.5 *P* = 0.0004, E12.5 versus E16.5 *P* = 0.0005, E14.5 versus E15.5 *P* = 0.0019, E14.5 versus E16.5 *P* < 0.0058; E11.5 n = 31, E12.5 n = 261; N = 146, E13.5 n = 43; N = 40, E14.5 n = 60; N = 48, E15.5 n = 16; N = 4, E16.5 n = 30; N = 9). (A3) Graph showing how the threshold of regenerative events (APs and plateau potentials) of V1^R^ changes with embryonic age. The average threshold became significantly more hyperpolarized after E12.5 (Kruskall-Wallis test *P* < 0.0001; E12.5 versus E11.5 *P* = 0.676, E12.5 versus E13.5 *P* = 0.0039, E12.5 versus E14.5 *P* < 0.0001, E12.5 versus E15.5 *P* < 0.0001, E12.5 versus E16.5 *P* < 0.0001, E13.5 versus E14.5 *P* > 0.999, E13.5 versus E15.5 *P* = 0.1398, E13.5 versus E16.5 *P* = 0.0013; E14.5 versus E15.5 *P* > 0.999, E14.5 versus E16.5 *P* = 0.0634, E15.5 versus E16.5 *P* > 0.999; E11.5 n = 20; N = 16, E12.5 n = 162; N = 139, E13.5 n = 31; N = 28, E14.5 n = 30; N = 26, E15.5 n = 16; N = 4, E16.5 n = 30; N = 9). Yellow and purple bars below the graphs indicate the two important phases of the functional development of spinal cord networks. The first one is characterized by synchronized neuronal activity (SNA) and the second one is characterized by the emergence of a locomotor-like activity (see text). Note that changes in *C*_*in*_ and *R*_*in*_ occurred at the end of the first developmental phase. (\**P* < 0.05, ** *P* < 0.01, *** *P* < 0.001; control, E12.5). The intrinsic activation properties were analyzed using 2 s suprathreshold depolarizing current steps. (B) Representative traces of voltage responses showing Single Spiking (SS) V1^R^ (gray), Repetitive Spiking (RS) V1^R^ (red), ME V1^R^ (purple) and Plateau Potential (PP) V1^R^ (blue) at E11.5 (B1), E13.5 (B2), E14.5 (B3) E15.5 (B4) and E16.5 (B5). (C) Bar graph showing how the proportions of the different firing patterns change from E11.5 to E16.5 (E11.5 n = 22; N = 18, E12.5 n = 163; N = 140, E13.5 n = 32; N = 29, E14.5 n = 57; N = 45, E15.5 n = 15; N = 4, E16.5 n = 28; N = 9). Yellow and purple bars below the graphs indicate the first and the second phase of functional embryonic spinal cord networks. The proportions of the different firing patterns significantly changed between E11.5 to E12.5 (Fisher’s exact test, *P* = 0.0052) with a significant increase in the proportion of RS V1^R^ (Fisher’s exact test, *P* = 0.0336) and a significant decrease in the proportion of ME V1^R^ (Fisher’s exact test, *P* = 0.01071) at E12.5. Only two firing patterns (SS and RS) were observed after E13.5 and most embryonic V1^R^ lost their ability to sustain tonic firing after E13.5. However, at E16.5 the proportion of RS V1^R^ significantly increased at the expense of SS V1^R^ when compared to E14.5 (Fisher’s exact test, *P* = 0.0112), indicating that embryonic V1^R^ began to recover the ability to sustain tonic firing after E15.5.

**Figure 5.**
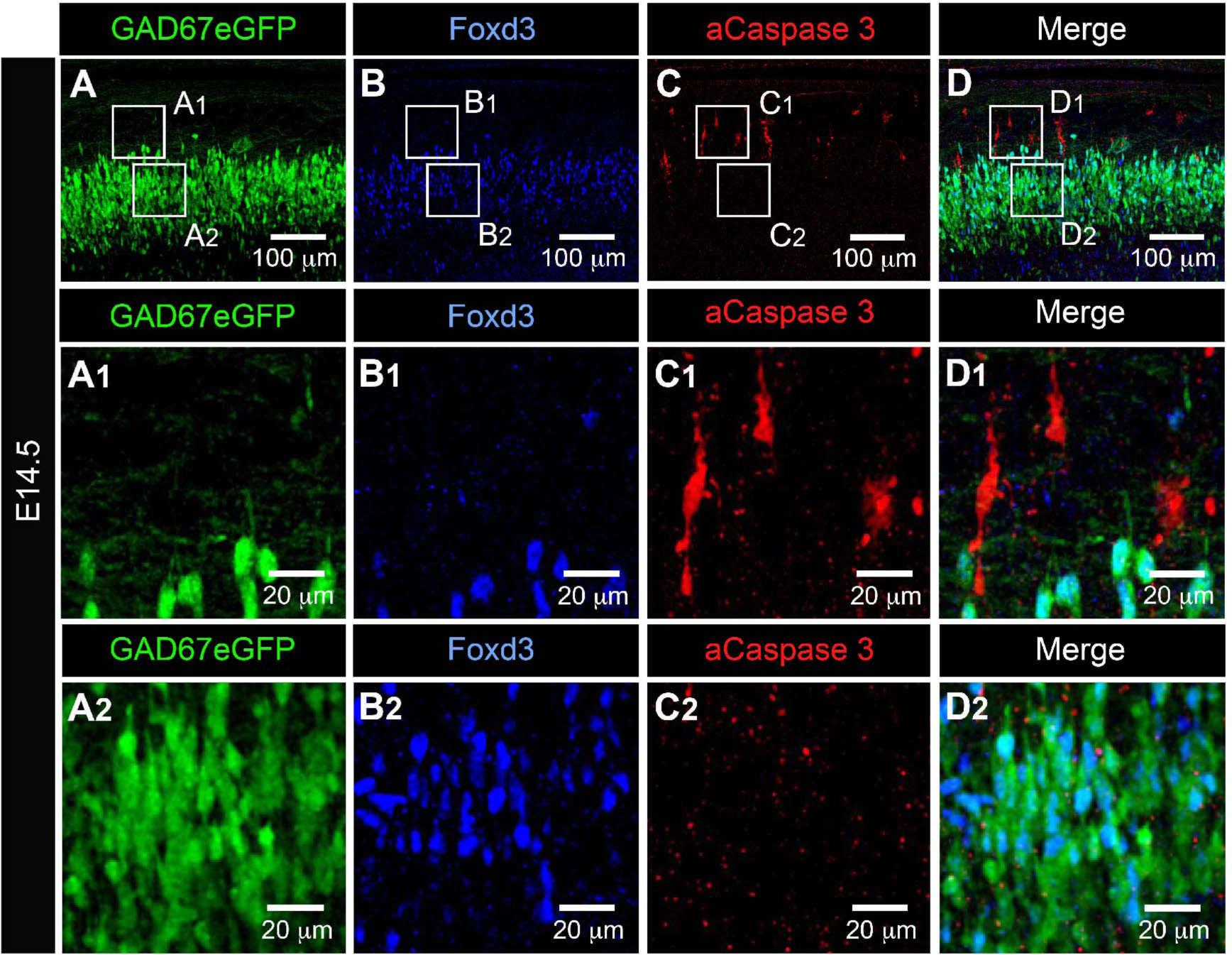
Activated caspase-3 is not observed in embryonic V1^R^ at E14.5. Representative confocal image of the ventral part of an isolated lumbar spinal cord of E14.5 GAD67-eGFP mouse embryo showing immunostainings using antibodies against eGFP (A), FoxD3 (B) and activated Caspase 3 (aCaspace 3) (C). (D) Superimposition of the three stainings shows that embryonic V1^R^ (eGFP+ and FoxD3+) were not aCaspase 3 immunoreactive. (A1, B1, C1 and D1). Enlarged images from A, B and C showing that aCaspase 3 staining is localized in areas where eGFP and Foxd3 staining were absent. (A2, B2, C2 and D2) Enlarged images from A, B and C showing that aCaspase 3 staining is absent in the area where V1^R^ (eGFP+ and FoxD3+) are located. aCaspase 3 staining that did not co-localize with GAD67eGFP likely indicates MN developmental cell death.

Spike threshold also decreased significantly between the first and the second developmental phases, dropping from about -34 mV at E12.5 to about -41 mV at E16.5 (***Figure 4A3***). Interestingly, this developmental transition around E14.5 correspond to the critical stage at which SNA gives way to a locomotor-like activity [11, 46, 47] and rhythmic activity becomes dominated by glutamate release rather than acetylcholine release [4].

This led us to hypothesize that this developmental transition could be also critical for the maturation of V1^R^ firing patterns. The distinct firing patterns observed at E12.5 were already present at E11.5 (***Figure 4B1,C***), but the percentage of RS V1^R^ strongly increased from E11.5 to E12.5, while the percentage of ME V1^R^ decreased significantly (***Figure 4C***). The heterogeneity of V1^R^ firing patterns then substantially diminished. Plateau potentials were no longer observed at E13.5 (***Figure 4B2,C***), and ME V1^R^ disappeared at E14.5 (***Figure 4B3,C***). Interestingly, the proportion of SS V1^R^ remained high from E13.5 to E15.5 and even slightly increased (91.23% at E14.5 and 93.33% at E15.5; ***Figure 4C***). This trend was partially reversed at E16.5, as the percentage of RS V1^R^ increased at the expense of SS V1^R^ (67.86% SS V1^R^ and 32.34% RS V1^R^; ***Figure 4B5,C***). This decrease in repetitive firing capability after E13.5 was surprising in view of what is classically admitted on the developmental pattern of neuronal excitability [18, 48]. Therefore, we verified that it did not reflect the death of some V1^R^ after E13.5. Our data did not reveal any activated caspase3 (aCaspase3) staining in V1^R^ (FoxD3 staining) at E14.5 (n = 10 SCs; N = 10) (***Figure 5***), in agreement with previous reports showing that developmental cell death of V1^R^ does not occur before birth [49].

To determine whether *G*_*Nap*_ and *G*_*Kdr*_ also controlled the firing pattern of V1^R^ at E14.5 (see ***Figure 4B3,C***), we assessed the presence of *I*_*Nap*_ and *I*_*Kdr*_ in single spiking V1^R^ at this embryonic age. Both *I*_*Nap*_ and *I*_*Kdr*_ were present in V1^R^ at E14.5 (***Figure 6––figure supplement 1* and *Figure 6––figure supplement 2*)** whereas, as in V1^R^ at E12.5, no calcium-dependent potassium current was detected at this developmental age (not shown). In SS V1^R^, *G*_*Kdr*_ was significantly higher at E14.5 (11.11 ± 1.12 nS, n = 10; N = 10) than at E12.5 (***Figure 1D***). In contrast, *G*_*Nap*_ was similar at E14.5 (0.13 ± 0.14 nS, n = 10; N = 10) and E12.5 **(*Figure 1E***). We also found that the *G*_*Nap*_/*G*_*Kdr*_ ratio was significantly lower for SS V1^R^ recorded at E14.5 (0.012 ± 0.004, n = 10) compared to RS V1^R^ (0.154 ± 0.022, n = 8) and PP V1^R^ (0.66 ± 0.132, n = 6) recorded at E12.5 (***Figure 1F***).

**Figure 6.**
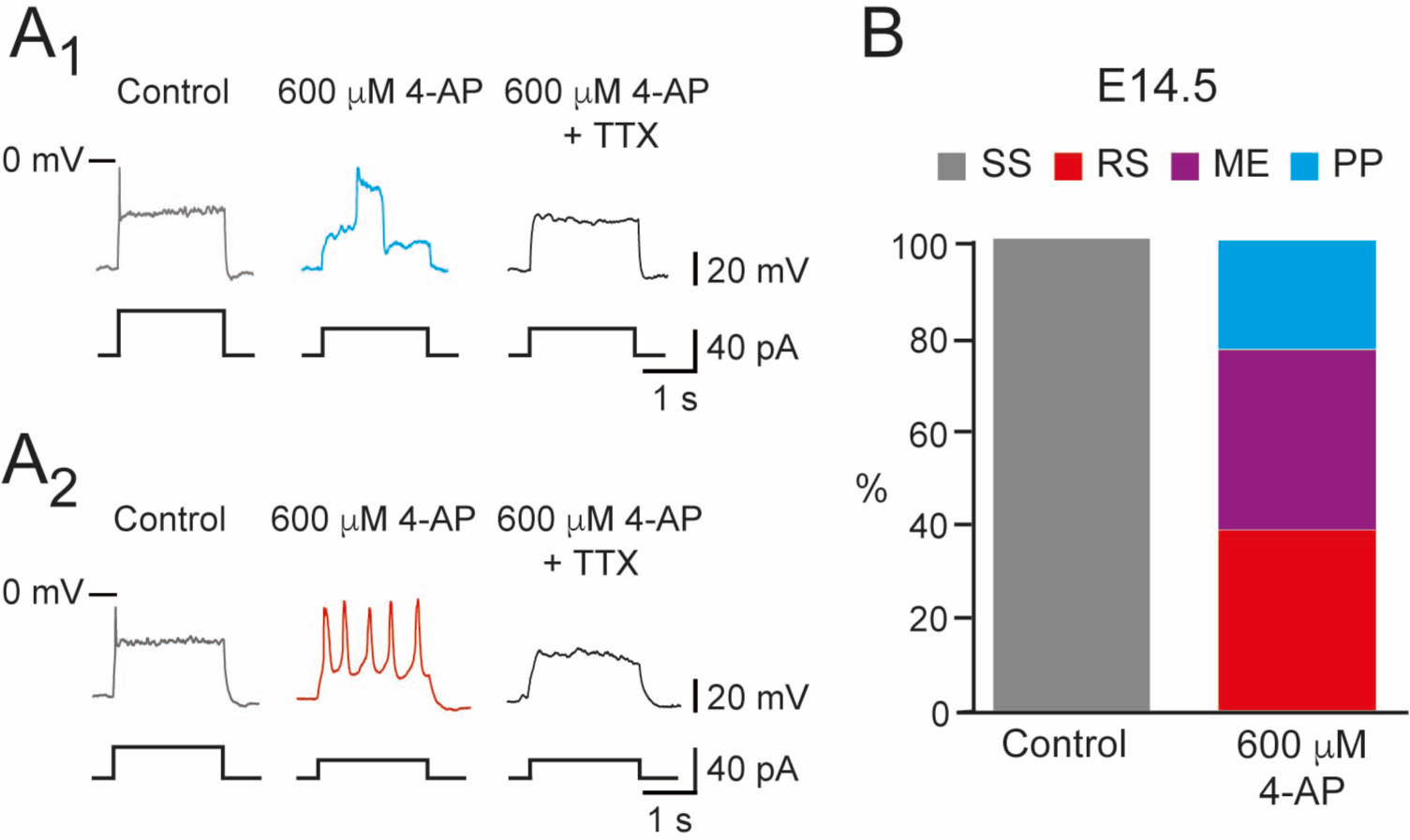
600 μM 4-AP changed the firing pattern of single spiking embryonic V1^R^ recorded at E14.5. The firing pattern of embryonic V1^R^ was evoked by 2 s suprathreshold depolarizing current steps. (A) Representative traces showing the effect of 4-AP application (600 μM) on the firing pattern of single spiking (SS) V1^R^ recorded at E14.5. Note that the applications of 600 μM 4-AP evoked either a plateau potential (A1) or repetitive spiking (A2), both fully blocked by TTX. (B) Bar plots showing the proportions of the different firing patterns observed in the presence of 600 μM 4-AP versus control recorded in SS V1^R^ at E14.5 (n = 14; N = 14). Single Spiking (SS) V1^R^ (gray), Repetitive Spiking (RS) V1^R^ (red), Mixed Events (ME) V1^R^ (purple), Plateau Potential (PP) V1^R^ (blue).

**Figure 7.**
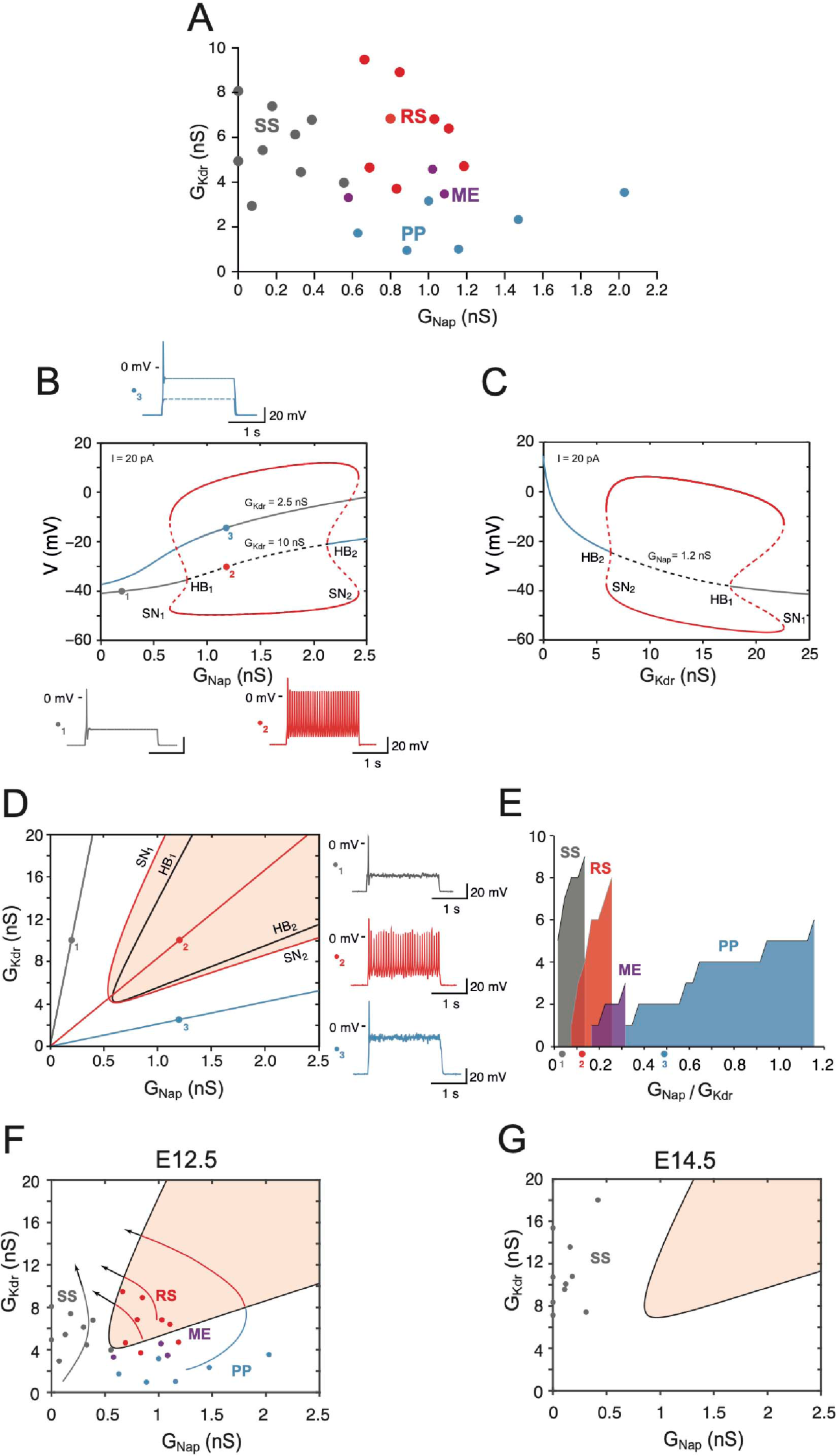
Embryonic V1^R^ firing patterns predicted by computational modeling. (A) Firing patterns of 27 recorded cells, in which both *G*_*Nap*_ and *G*_*Kdr*_ were measured. Gray: SS, red: RS, blue: PP. The three purple points located at the boundary between the RS and PP regions correspond to mixed events (ME) where plateau potentials alternate with spiking episodes. Note that no cell exhibited low values of both *G*_*Nap*_ and *G*_*Kdr*_(lower left), or large values of both conductances (upper right). (B) Bifurcation diagram of the deterministic model when *G*_*Kdr*_ is kept fixed to 2.5 nS or 10 nS while *G*_*Kdr*_ is varied between 0 and 2.5 nS. *G*_*in*_= 1 nS and *I* = 20 pA. For *G*_*Kdr*_= 10 nS (i.e., in the top experimental range), the red curves indicate the maximal and minimal voltages achieved on the stable limit cycle associated with repetitive firing (solid lines) and on the unstable limit cycle (dashed lines). The fixed point of the model is indicated by a gray solid line when it corresponds to the stable quiescent state, a gray dashed line when it is unstable and a solid blue line when it corresponds to a stable plateau potential. The two HB corresponding to the change of stability of the quiescence state (HB_1_, *G*_*Nap*_= 0.81 nS) and of the voltage plateau (HB_2_, *G*_*Nap*_= 2.13 nS) are indicated, as well as the two SN bifurcations of limit cycles associated with the onset (SN_1_, *G*_*Nap*_ = 0.65 nS) and offset (SN_2_, *G*_*Nap*_= 2.42 nS) of repetitive spiking as *G*_*Nap*_ is increased. For *G*_*Kdr*_= 2.5 nS, the model does not display repetitive firing; it possesses a unique fixed point, which is always stable (blue-gray curve). The transition from quiescence to plateau is gradual with no intervening bifurcation. Representative voltage traces of the three different activity patterns are shown: single spiking in response to a 2 s current pulse (gray, *G*_*Nap*_= 0.2 nS, *G*_*Kdr*_= 10 nS), repetitive spiking (red, *G*_*Nap*_= 1.2 nS, *G*_*Kdr*_= 10 nS) and plateau potential (blue, *G*_*Nap*_= 1.2 nS, *G*_*Kdr*_= 2.5 nS). Note that the plateau never outlasts the current pulse. (C) Bifurcation diagram when *G*_*Nap*_ is kept fixed at 1.2 nS and *G*_*Kdr*_ is varied between 0 and 25 nS (*I* = 20 pA). Same conventions as in B. Plateau potential is stable until the subcritical Hopf bifurcation HB_2_ (*G*_*Kdr*_= 6.34 nS) is reached, repetitive firing can be observed between SN_2_ (*G*_*Kdr*_= 5.93 nS) and SN_1_ (*G*_*Kdr*_= 22.65 nS). The quiescent state is stable from point HB_1_ (*G*_*Kdr*_= 17.59 nS) onward. (D) Two-parameters bifurcation diagram of the model in the *G*_*Kdr*_-*G*_*Kdr*_ plane (*I* = 20 pA). The black merged curves indicate the bifurcations HB_1_ and HB_2_. The red curves indicate the SN bifurcations of limit cycles SN_1_ and SN_2_. The shaded area indicates the region where repetitive firing can occur. The oblique lines through the points labeled 1, 2 and 3, the same as in B, correspond to three different values of the ratio *G*_*Kdr*_/*G*_*Kdr*_: 0.02 (gray), 0.12 (red) and 0.48 (blue). Voltage traces on the right display the response to a 2 s current pulse when channel noise is taken into account for the three regimes: quiescence (top, gray trace and dot in the diagram), repetitive firing (middle, red) and plateau potential (bottom, blue). They correspond to the three deterministic voltage traces shown in B. Note that the one-parameter bifurcation diagrams shown in B correspond to horizontal lines through points 1 and 2 (*G*_*Kdr*_= 10 nS) and through point 3 (*G*_*Kdr*_= 2.5 nS), respectively. The bifurcation diagram in C corresponds to a vertical line through point 2 and 3 (*G*_*Nap*_= 1.2 nS). (E) Cumulative distribution function of the ratio *G*_*Nap*_/*G*_*Kdr*_ for the four clusters in A, showing the sequencing SS (gray) à RS (red) à ME (purple, 3 cells only) à PP (blue) predicted by the two-parameters bifurcation diagram in D. The wide PP range, as compared to SS and RS, merely comes from the fact that *G*_*Kdr*_ is small for cells in this cluster. The three colored points indicate the slopes of the oblique lines displayed in D (0.02, 0.12 and 0.48, respectively). (F) The data points in A are superimposed on the two-parameters bifurcation diagram shown in D, demonstrating a good agreement between our basic model and experimental data (the same color code as in A for the different clusters). The bifurcation diagram is simplified compared to A, only the region where repetitive spiking is possible (i.e. between the lines SN_1_ and SN_2_ in A) being displayed (shaded area). Notice that 3 ME cells (purple dots) are located close to the transition between the RS and PP regions. The four arrows indicate the presumable evolution of *G*_*Nap*_ and *G*_*Kdr*_ for SS, RS, ME and PP cells between E12.5 and E14.5-15.5. *G*_*Nap*_ eventually decreases while *G*_*Kdr*_ keeps on increasing. (G) Distribution of a sample of cells in the *G*_*Kdr*_-*G*_*Kdr*_ plane at E14.5. All the cells are located well within the SS region far from bifurcation lines because of the decreased *G*_*Nap*_ compared to E12.5, the increased *G*_*Kdr*_, and the shift of the RS region (shaded) due to capacitance increase (18 versus 13 pF).

We tested the effect of 4-AP in SS V1^R^ at E14.5. At this embryonic age, 300 μM 4-AP inhibited only 59.2% of *I*_*Kdr*_. Increasing 4-AP concentration to 600 μM did not inhibit *I*_*Kdr*_ significantly more (60.2%) (***Figure 6––figure supplement 2***), indicating that inhibition of *I*_*Kdr*_ by 4-AP reached a plateau at around 300 μM. 600 μM 4-AP application had no significant effect on *I*_*A*_ (***Figure 6––figure supplement 2***). The application of the maximal concentration of 4-AP tested (600 μM) converted SS V1^R^ (n = 13; N = 13) to PP V1^R^ (23.1%; ***Figure 6A1,B***), RS V1^R^ (38.5%; ***Figure 6A2,B***) or ME V1^R^ (38.4%; ***Figure 6B***), as was observed at E12.5, thus indicating that the firing pattern of V1^R^ depends on the balance between *I*_*Nap*_ and *I*_*Kdr*_ also at E14.5. Plateau potential and repetitive spiking recorded in the presence of 4-AP at E14.5 were fully blocked by 0.5-1 μM TTX indicating that they were generated by voltage-gated sodium channels (***Figure 6A1,A2***), as observed at E12.5.

### Theoretical analysis: the basic model

As shown in ***Figure 7A*** for 26 cells, in which both *G*_*Nap*_ and *G*_*Kdr*_ were measured, the three largest clusters revealed by the hierarchical clustering analysis (SS, RS and PP, which account together for the discharge of more than 95% of cells, see ***Figure 5***) correspond to well defined regions of the *G*_*Nap*_ - *G*_*Kdr*_ plane. Single spiking is observed only when *G*_*Kdr*_ is smaller than 0.6 nS. For larger values of *G*_*Nap*_, repetitive spiking occurs when *G*_*Kdr*_ is larger than 3.5 nS, and V1^R^ display plateau potentials when *G*_*Kdr*_ is smaller than 3.5 nS. Mixed events (ME, 4.5% of the 163 cells used in the cluster analysis), where plateaus and spiking episodes alternate, are observed at the boundary of RS and PP clusters. This suggested to us that a conductance-based model incorporating only the leak current, *I*_*Nat*_, *I*_*Nap*_ and *I*_*Kdr*_ (see Materials and Methods) could account for most experimental observations, the observed zonation being explained in terms of bifurcations between the different stable states of the model. Therefore, we first investigated a simplified version of the model without *I*_*A*_ and slow inactivation of *I*_*Nap*_.

A one-parameter bifurcation diagram of this “basic” model is shown in ***Figure 7B*** for two values of *G*_*Kdr*_(2.5 nS and 10 nS) and a constant injected current *I* = 20 pA. In both cases, the steady-state membrane voltage (stable or unstable) and the peak and trough voltages of stable and unstable periodic solutions are shown as the function of the maximal conductance *G*_*Nap*_ of the *I*_*Nap*_ current, all other parameters being kept constant. For *G*_*Kdr*_= 10 nS, the steady-state membrane voltage progressively increases (in gray) with *G*_*Nap*_, but repetitive spiking (in red, see voltage trace for *G*_*Nap*_= 1.5 nS) is not achieved until *G*_*Nap*_ reaches point SN_1_, where a saddle node (SN) bifurcation of limit cycles occurs. This fits with the experimental data, where a minimal value of *I*_*A*_ is required for repetitive spiking (see also [25], and is in agreement with the known role of *I*_*Kdr*_ in promoting repetitive discharge [50, 51]. Below SN_1,_ the model responds to the onset of a current pulse by firing only one spike before returning to quiescence (see voltage trace for *G*_*Nap*_= 0.2 nS), or a few spikes when close to SN_1_ (not shown) before returning to quiescence. The quiescent state becomes unstable through a subcritical Hopf bifurcation (HB) at point HB_1_, with bistability between quiescence and spiking occurring between SN_1_ and HB_1_ points. Repetitive firing persists when *G*_*Nap*_ is increased further and eventually disappears at point SN_2_. The firing rate does not increase much throughout the RS range (***Figure 7–figure supplement 1C***), remaining between 20.1 Hz (at SN_1_) and 28.7 Hz (at SN_2_). A stable plateau appears at point HB_2_ through a subcritical HB. The model is bistable between HB_2_ amplitude APs coexisting in this range. and SN_2_, with plateau and large

The model behaves very differently when *G*_*Kdr*_ is reduced to 2.5 nS (gray-blue curve in ***Figure 7B***). It exhibits a unique stable fixed point whatever the value of *G*_*Nap*_ is, and the transition from quiescence to plateau is gradual as *G*_*Nap*_ is increased. No repetitive spiking is ever observed. This indicates that the activity pattern is controlled not only by *G*_*Nap*_ but also by *G*_*Kdr*_. This is demonstrated further in ***Figure 7C***, where *G*_*Kdr*_ was fixed at 1.2 nS while *G*_*Kdr*_ was increased from 0 to 25 nS. The model exhibits a plateau potential until *G*_*Kdr*_ is increased past point the subcritical HB point HB_2_, repetitive spiking sets in before at point SN_2_ *via* a SN of limit cycles bifurcation. When *G*_*Kdr*_ is further increased, repetitive firing eventually disappears through a SN bifurcation of limit cycles at point SN_1_, the quiescent state becomes stable through a subcritical HB at point HB_1_, and bistability occurs between these two points. This behavior is in agreement with ***Figure 7A***.

Since both conductances *G*_*Nap*_ and *G*_*Kdr*_ control the firing pattern of embryonic V1^R^ cells, we computed a two-parameters bifurcation diagram (***Figure 7D***), where the stability regions of the different possible activity states and the transition lines between them are plotted in the *G*_*Nap*_-*G*_*Kdr*_ plane. The black curves correspond to the bifurcations HB_1_ and HB_2_ and delimit a region where only repetitive firing occurs. The red curves correspond to the SN bifurcations of periodic orbits associated with the transition from quiescence to firing (SN_1_) and the transition from plateau to firing (SN_2_). They encompass a region (shaded area) where repetitive firing can be achieved but may coexist with quiescence (between the HB_1_ and SN_1_ lines) or plateau potential (in the narrow region between the HB_2_ and SN_2_ lines).

Some important features of the diagram must be emphasized: 1) minimal values of both *G*_*Nap*_ (to ensure sufficient excitability) and *G*_*Kdr*_(to ensure proper spike repolarization) are required for repetitive spiking, 2) quiescence and plateau can be clearly distinguished only when they are separated by a region of repetitive spiking (see also ***Figure 7B*** for *G*_*Kdr*_= 10 nS), otherwise the transition is gradual (***Figure 7B*** for *G*_*Kdr*_= 2.5 nS), 3) only oblique lines with an intermediate slope cross the bifurcation curve and enter the RS region (see, for example, the red line in ***Figure 7D***). This means that repetitive spiking requires an appropriate balance between *I*_*Nap*_ and *I*_*Kdr*_. If the ratio *G*_*Nap*_/*G*_*Kdr*_ is too large (blue line) or too small (gray line), only plateau potentials or quiescence will be observed at steady state. This is exactly what is observed in experiments, as shown by the cumulative distribution function of the ratio *G*_*Nap*_/*G*_*Kdr*_ for the different clusters of embryonic V1^R^ in ***Figure 7E*** (same cells as in ***Figure 7A***). The ratio increases according to the sequence SS → RS → ME → PP, with an overlap of the distributions for SS V1^R^ and RS V1^R^. Note also that the ratio for ME cells (around 0.25) corresponds to the transition between repetitive spiking and plateau potentials (more on this below).

Embryonic V1^R^ cells display voltage fluctuations that may exceed 5 mV and are presumably due to channel noise. The relatively low number of sodium and potassium channels (of the order of a few thousands) led to voltage fluctuations in the stochastic version of our model comparable to those seen experimentally when the cell was quiescent (top voltage trace in ***Figure 7D***) or when a voltage plateau occurred (bottom trace). Channel noise caused some jitter during repetitive spiking (middle trace), and induced clearly visible variations in the amplitude of APs. However, repetitive firing proved to be very robust and was not disrupted by voltage fluctuations. Altogether, channel noise little alters the dynamics (compare the deterministic voltage traces in ***Figure 7B*** and the noisy traces in ***Figure 7D***). This is likely because channel noise has a broad power spectrum and displays no resonance with the deterministic solutions of the model.

The one-parameter bifurcation diagram of the model was not substantially modified when we took *I*_*A*_ into account, as shown in ***Figure 6––figure supplement 1***. It just elicited a slight membrane hyperpolarization, an increase in the minimal value of *G*_*Nap*_ required for firing, and a decrease of the firing frequency. The transition from repetitive firing to plateau was not affected because *I*_*A*_ is then inactivated by depolarization.

The bifurcation diagram of ***Figure 7D*** accounts *qualitatively* for the physiological data on V1^R^ at E12.5 presented in ***Figure 7A***, as shown in ***Figure 7F*** where the conductance data of ***Figure 7A*** were superimposed on it. However, one must beware of making a more *quantitative* comparison because the theoretical bifurcation diagram was established for a constant injected current of 20 pA, whereas the current injected in experiments data varied from neuron to neuron and ranged from 10 to 30 pA in the sample shown in ***Figure 7A***. The position of bifurcation lines in the *G*_*Nap*_-*G*_*Kdr*_ plane depends not only on the value of the injected current, but on the values chosen for the other parameters, which also vary from cell to cell but were kept at fixed values in the model (Ori et al. 2018). For instance, the diagrams were computed in ***Figure 7D,F*** for *G*_*in*_= 1 nS and *C*_*in*_= 13 pF, the median values of the input conductance and capacitance at E12.5, taking no account of the cell-to-cell variations of these quantities. Between E12.5 and E14.5, *C*_*in*_ which provides an estimate of the cell size, increases by 38% in average, whereas *G*_*in*_ is not significantly modified (see ***Figure 4***). As illustrated in ***Figure 7G*** the two-parameters bifurcation diagram is then shifted upward and rightward compared to ***Figure 7F***, because larger conductances are required to obtain the same firing pattern. The observed regression of excitability from E12.5 to E14.5- E15.5 (see ***Figure 4C***) thus comes from a decrease in *G*_*Nap*_ density (see presumable developmental trajectories indicated by arrows in ***Figure 7F***) together with a shift of the RS region as cell size increases. As a result, all 10 cells shown in ***Figure 7G*** are deeply inside the SS region at E14.5.

It is less straightforward to explain on the basis of our model the experiments where 4-AP changed the firing pattern of SS V1^R^ (***Figure 2***). Indeed, the decrease of *G*_*Kdr*_(***Figure 6–– figure supplement 2***), although it may exceed 70% at the higher concentrations of 4-AP we used, is not sufficient by itself to account for the change in the firing pattern of V1^R^ (***Figure 6––figure supplement 2***) because data points in the SS cluster will not cross the bifurcation lines between SS and RS (SN1) and between RS and PP (SN2) when displaced downward in the *G*_*Kdr*_- *G*_*Nap*_ plane. However, 4-AP at a 300 μM concentration also decreases *G*_*in*_(by 23% in average and up to 50% in some neurons), the rheobase current with it, and the current that was injected in cells during experiments was reduced accordingly. hen we take into account this reduction of both *G*_*in*_ and *I* the two parameters bifurcation diagram pf the model remains qualitatively the same, but it is shifted leftward and downward in the *G*_*Nap*_- *G*_*Kdr*_ plane (***Figure 6––figure supplement 2***). As a consequence, the bifurcation lines between SS and RS and between RS and PP (SN2) are then successively crossed when *G*_*Kdr*_ is reduced, in accordance with experimental results.

### Theoretical analysis: slow inactivation of I_Nap_ and bursting

Our basic model accounts for the firing pattern of 73% of the 163 cells used in the cluster analysis. However, bursting, under the form of recurring plateaus separated by brief repolarization episodes (see a typical trace in ***Figure 8A left***), was experimentally observed in half of PP V1^R^ (24 out of 46), and plateaus intertwined with spiking episodes were recorded in the 13 cells of the ME cluster (8% of the total sample, see ***Figure 8A right*** for a typical example). Recurrent plateaus indicate membrane bistability and require that the *I* − *V* curve be S-shaped. This occurs when *G*_*Nap*_ is large and *G*_*Kdr*_ small. (***Figure 8B1,B2***). However, our basic model lacks a mechanism for switching between quiescent state and plateau, even in this case. Channel noise might induce such transitions, but our numerical simulations showed that this is too infrequent to account for bursting (see voltage trace in ***Figure 8B1*** where the plateau state is maintained despite channel noise).

**Figure 8.**
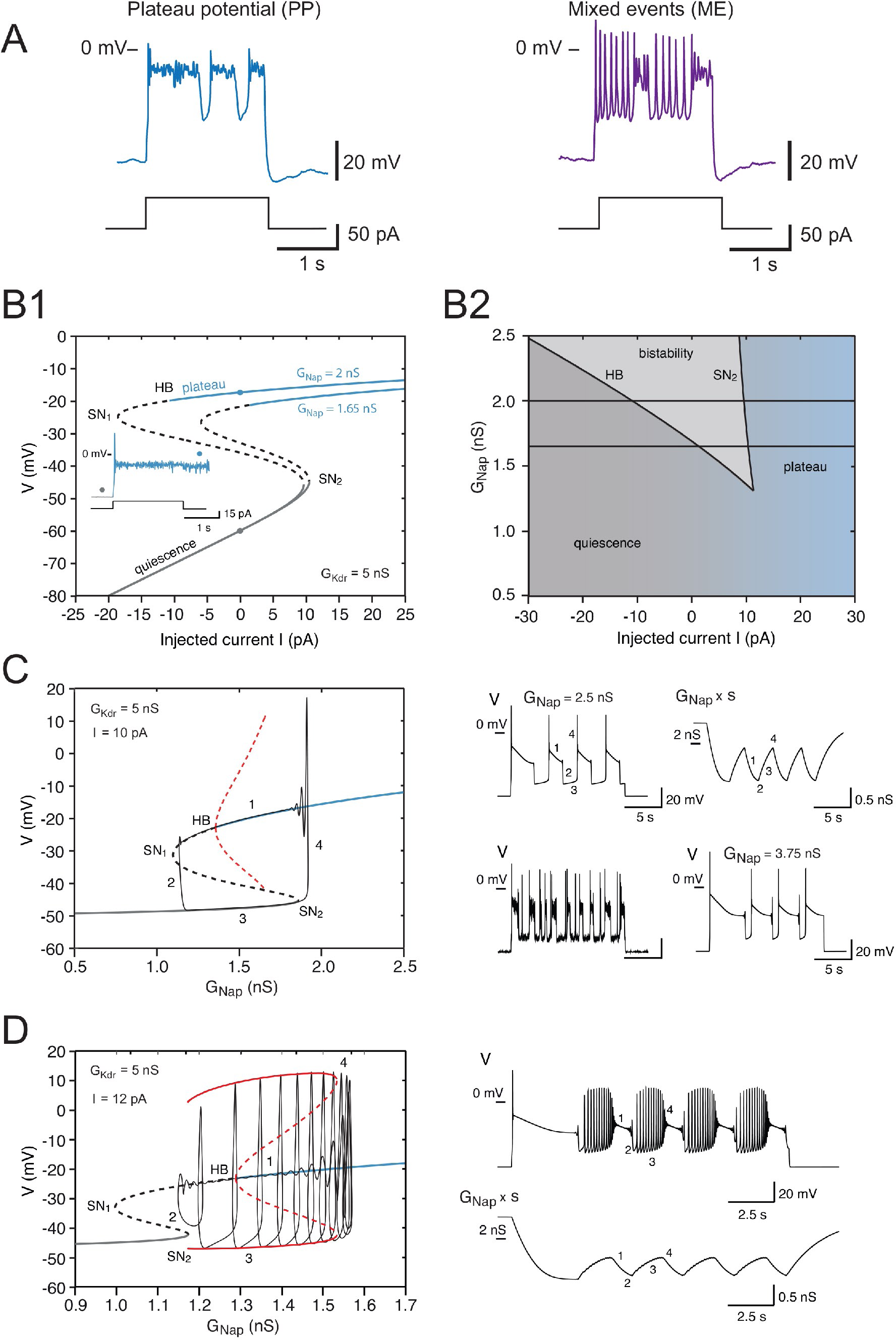
Effects of the slow inactivation of *I*_*Nap*_ on firing patterns predicted by computational modeling. (A) Examples of repetitive plateaus (left) and mixed events (right) recorded in V1^R^ at E12.5 during a 2 s current pulse. (B1) Current-voltage curve of the basic model (without slow inactivation of *I*_*Nap*_, and without *I*_*A*_ or channel noise) for *G*_*Kdr*_= 5 nS and for *G*_*Nap*_= 1.65 nS (lower curve) and 2 nS (upper curve). Solid lines denote stable fixed points and dashed lines unstable ones. For *G*_*Nap*_=1.65 nS, bistability between quiescence and plateau occurs between 1.39 and 10.48 pA. When *G*_*Nap*_ is increased to 2 nS, the bistability region ranges from -10.84 to 9.70 pA, thus extending into the negative current range. This implies that once a plateau has been elicited, the model will stay in that stable state and not return to the resting state, even though current injection is switched off (see insert). B1 Insert. Voltage response to a 2 s current pulse of 15 pA for *G*_*Nap*_= 2 nS. The resting state (gray dot on the lower curve in B1 is destabilized at pulse onset and a plateau is elicited (blue dot on the upper curve in B1). At pulse offset, the plateau is maintained, even though the injected current is brought back to zero, and channel noise is not sufficient to go back to the resting state. (B2) Domain of bistability between quiescence and plateau (shaded) in the *I* - *G*_*Nap*_ plane for *G*_*Kdr*_= 5 nS. It is delimited by the line SN_2_ where a SN bifurcation of fixed points occurs and by the subcritical Hopf bifurcation line HB where the plateau becomes unstable. Bistability requires that *G*_*Nap*_ exceeds 1.35 nS, and the domain of bistability enlarges as *G*_*Nap*_ is increased further. The two horizontal lines correspond to the two cases shown in B1: *G*_*Nap*_ =1.65 nS and 2 nS. (C) Behavior of the model when slow inactivation is incorporated. The bifurcation diagram of the basic model (without slow inactivation) for *I* = 10 pA and *G*_*Kdr*_= 5 nS (same conventions as in Fig 7B) and the stable limit cycle (black solid curve) obtained when slow inactivation is added are superimposed. The limit cycle is comprised of four successive phases (see labels): 1) long plateau during which *I*_*Nap*_ slowly inactivates, 2) fast transition to the quiescent state, 3) repolarization episode during which *I*_*Nap*_ slowly deinactivates, 4) fast transition back to the plateau. Each plateau starts with a full-blown action potential followed by rapidly decaying spikelets. Note that the bifurcation HB is subcritical here (unstable limit cycle shown by dashed red curve), at variance with square wave bursting (supercritical bifurcation and stable limit cycle); this is a characteristic feature of pseudo-plateau bursting. Note also that the plateau extends beyond the bifurcation HB because it is only weakly unstable then. Responses to a 15 s current pulse are shown on the right side. Top left: voltage response (*G*_*Nap*_= 2.5 nS), Top right: behavior of the “effective” conductance of *I*_*Nap*_ channel, i.e., the maximal conductance *G*_*Nap*_ multiplied by the slow inactivation variable *s*. Bottom left: voltage trace when channel noise is added to fast and slow gating variables, Bottom right: Voltage trace when *G*_*Nap*_ is increased by 50% to 3.75 nS. (D) Mixed events. The bifurcation diagram of the basic model for *G*_*Kdr*_= 5 nS and *I* = 12 pA and the stable limit cycle obtained in the presence of slow inactivation (*G*_*Nap*_= 2.5 nS) are superimposed. Here again, the limit cycle is comprised of four successive phases (see labels): 1) slow inactivation of *I*_*Nap*_ that leads to the crossing of the bifurcation point HB_2_ and then to the destabilization of the plateau potential, 2) fast transition to the spiking regime, 3) repetitive spiking during which *I*_*Nap*_ slowly de-inactivates, which leads to the crossing of the bifurcation point SN_2_ and terminates the spiking episode, 4) fast transition back to the stable plateau potential. Response to a 15 s current pulse of 12 pA is shown on the right in the absence of any channel noise. Top: Voltage trace (same labels as in the bifurcation diagram on the left), Bottom: Variations of the “effective” conductance *G*_*Nap*_*s* (same labels as in the voltage trace). Note that de-inactivation sufficient to trigger a new plateau occurs over a series of successive spikes, hence the small oscillations are visible on the trace. Note also that in C and D the first plateau lasts longer than the following ones, as in electrophysiological recordings of embryonic V1^R^ cells displaying repetitive plateaus. This form of adaptation is caused by the slow inactivation of the persistent sodium current.

To explain recurrent plateaus during a constant current pulse, we have to incorporate in our model an additional slow dynamical process. Therefore, we took into account the slow inactivation of *I*_*Nap*_ that is observed in experiments. *I*_*Kdr*_ also inactivates slowly but over times that are much longer than the timescale of bursting, which is why we did not take its slow inactivation into account. The one-parameter bifurcation diagram of the basic model without slow inactivation of *I*_*Nap*_ is shown in ***Figure 8C*** for *G*_*Kdr*_= 5 nS and an injected current reduced to 10 pA (as compared to 20 pA in the previous section), so as to allow for bistability (see ***Figure 8B2***). The *G*_*Nap*_− *V* curve is then S-shaped, as shown in ***Figure 8B1***, with a bistability region for *G*_*Nap*_ between 1.36 and 1.85 nS. This is in contrast with ***Figure 7B*** where the *G*_*Kdr*_ − *V* curve was monotonic. Adding the slow (de)inactivation of *I*_*Nap*_ then causes periodic transitions between up (plateau) and down (quiescent) states, as illustrated by the top voltage trace on the right of ***Figure 8C***, and the model displayed a stable limit cycle (shown in black in the bifurcation diagram on the left of ***Figure 8C***). This mechanism is known as pseudo-plateau or plateau-like bursting (a.k.a. fold-subcritical HB bursting) [52]. It is akin to square wave bursting [53-56], but the up-state is a stable fixed point rather than a limit cycle [57-59], which is why recurrent plateaus are obtained rather than bursts of APs. The duration of the plateaus and repolarization episodes depends on the values of *G*_*Nap*_ and *G*_*Kdr*_. A voltage-independent time constant τ_*s*_= *2 ms* leads to up and down states of comparable durations (see top left voltage trace in ***Figure 8C***). In agreement with the bifurcation diagram of ***Figure 8C***, the persistent sodium current inactivates during plateaus (phase 1, see bottom right trace in in ***Figure 8C***) and de-inactivates during quiescent episodes (phase 3, see bottom right trace). Transitions from the down-state to the up-state occurs when inactivation has reached its maximal value (phase 2) and transition from the up-state to the down state when it has reached its maximum (phase 4). Adding channel noise preserves bursting but introduces substantial randomness in the duration of plateaus and repolarization episodes (bottom left voltage trace in ***Figure 8C***). Moreover, it substantially decreases the duration of both plateaus and quiescent episodes by making transition between the two states easier (compare the top and bottom voltage traces on the left, both computed for τ_*s*_ = *2 ms*).

Increasing *G*_*Nap*_(or decreasing *G*_*Kdr*_) makes plateaus much longer than quiescent episodes (see bottom right voltage trace in ***Figure 8C***). This again points out to the fact that the ratio is an important control parameter. We also noted that adding the *I*_*A*_ current lengthened the quiescence episodes (***Figure 6––figure supplement 1***).

Slow inactivation of *I*_*Nap*_ also provides an explanation for mixed patterns, where plateaus alternate with spiking episodes (***Figure 8A, right***). They take place in our model near the transition between repetitive spiking and plateau, as in experiments (see ***Figure 8A***). Slow inactivation can lead to elliptic bursting, notably when the bifurcation HB_2_ is subcritical [60, 61], which is the case here (***Figure 8D***). The model then displays a stable limit cycle with alternating plateaus and spiking episodes, arising from crossing the bifurcation points HB_2_ and SN_2_ back and forth (see bifurcation diagram in ***Figure 8D*** and top voltage trace). We note that sufficient de-inactivation of *I*_*Nap*_ for triggering a new plateau (phase 3 in the bottom trace of ***Figure 8D***) may be difficult to achieve during spiking episodes, because voltage oscillates over a large range, which tends to average out the variations of the inactivation level. If de-inactivation is not sufficient, the model keeps on spiking repetitively without returning to the plateau state. This is what occurs for cells well within the RS region, far away from the RS-PP transition. It also probably explains why it was difficult in many recorded cells to elicit plateaus by increasing the injected current, inactivation balancing then the increase of *I*_*Nap*_ induced by the larger current.

Altogether, our study shows that a model incorporating the slow inactivation of *I*_*Nap*_ accounts for all the firing patterns displayed by cells from the PP and ME clusters.

## Discussion

V1^R^ constitute a homogeneous population when referring to their transcription factor program during development [24, 62], their physiological function [63] and their firing pattern at postnatal stages [27]. Surprisingly, our electrophysiological recordings and our cluster analysis clearly indicate that distinct functional classes of V1^R^ are transiently present during development at the onset of the SNA (E11.5-E12.5). Five different groups of embryonic V1^R^ were defined using cluster analysis, according to their firing properties.

### Development of the firing pattern of embryonic V1^R^ during SNA

It is generally assumed that, during early development, newborn neurons cannot sustain repetitive firing [35, 48]. Later on, neurons progressively acquire the ability to fire repetitively, APs become sharper, and neurons eventually reach their mature firing pattern, due to the progressive appearance of a panoply of voltage-gated channels with different kinetics [18, 35, 48]. Our results challenge the general view that single spiking is a more primitive form of excitability [35]. Indeed, we show that repetitive firing and plateau potentials dominated at early stages (E11.5-E12.5), while single spiking was prevailing only later (E13.5- E16.5).

The different V1^R^ firing patterns observed at E11.5-E12.5 might reflect variability in the maturation level between V1^R^ at a given developmental stage, as suggested for developing MNs [64, 65]. However, this is unlikely since V1^R^ transiently lose their ability to sustain tonic firing or plateau potential after E13.5. The heterogeneous discharge patterns of V1^R^ observed before E13.5 contrasts with the unique firing pattern of V1^R^ at postnatal age [27]. Accordingly, the transient functional heterogeneity of V1^R^ rather reflects an early initial developmental stage (E11.5-E13.5) of intrinsic excitability.

The physiological meaning of the transient functional involution of V1^R^ that follows, after E12.5, is puzzling. To our knowledge, such a phenomenon was never described in vertebrates during CNS development. So far, a functional involution was described only for inner hair cells between E16 and P12 [66, 67] and cultured oligodendrocytes [68], and it was irreversible. Because most V1^R^ cannot sustain tonic firing after E12.5, it is likely that their participation to SNA is limited to the developmental period before other GABAergic interneuron subtypes mature and start to produce GABA and glycine [69]. Interestingly, embryonic V1^R^ begin to recover their capability to sustain tonic firing when locomotor-like activity emerges [4, 11], a few days before they form their recurrent synaptic loop with MNs (around E18.5 in the mouse embryos, [70]). One possible function of the transient involution between E12.5 and E15.5 could be to regulate the growth of V1^R^ axons toward their targets. It is indeed known that low calcium fluctuations within growth cones are required for axon growth while high calcium fluctuations stop axon growth and promote growth cone differentiation [71].

### Ion channels mechanisms underlying the functional heterogeneity of embryonic V1^R^

Blockade of *I*_*Nap*_ leads to single spiking [25], which emphasizes the importance of this current for the occurrence of repetitive firing and plateau potentials in V1^R^ at early developmental stages. But these neurons can also switch from one firing pattern to another, when *G*_*Kdr*_ is decreased by 4-AP, which emphasizes the importance of *I*_*Kdr*_. We found that the main determinant of embryonic V1^R^ firing pattern is the balance between *G*_*Nap*_ and *G*_*Kdr*_. A Hodgkin-Huxley-type model incorporating a persistent sodium current *I*_*Nap*_ provided a parsimonious explanation of all the firing patterns recorded in the V1^R^ population at E12.5. It provided a mathematical interpretation for the clustering of embryonic V1^R^ shown by the hierarchical analysis and accounted for the effect of 4-AP and riluzole [25] on the discharge. Remarkably, it highlighted how a simple mechanism involving only the two opposing currents *I*_*Nap*_ and *I*_*Kdr*_, but not *I_A_*, could produce functional diversity in a population of developing neurons. The model explained why minimal *G*_*Kdr*_ and *G*_*Kdr*_ are required for firing, and how a synergy between *G*_*Nap*_ and *G*_*Kdr*_ controls the firing pattern and accounts for the zonation of the *G*_*Nap*_ − *G*_*Kdr*_ plane that is observed experimentally.

Taking into account the slow inactivation of *I*_*Nap*_ to the model allowed us to explain the bursting patterns displayed by cells of the PP and ME clusters. We showed, in particular, that mixed events arose from elliptic bursting at the repetitive spiking-plateau transition and that smooth repetitive plateaus could be explained by a pseudo-plateau bursting mechanism [52, 58]. Such bursting scenario has been previously studied in models of endocrine cells [57, 72, 73] and adult neurons [74], but rarely observed in experiments.

Heterogeneity of the discharge pattern of pacemaker neurons has also been observed in the embryonic pre-Bötzinger network [75]. However, it was related there to the gradual change of balance between two inward currents, *I*_*Nap*_ and the calcium-activated nonselective cationic current *I*_*CAN*_ during neuronal maturation, which led to the progressive replacement of pseudo-plateau bursting by elliptic bursting. Such a scenario cannot account for the variety of discharge patterns observed in embryonic V1^R^ at the E11.5-12.5 stage of development [25]. Our theoretical analysis and our experimental data clearly indicate that the interplay between two opposing currents is necessary to explain all the firing patterns of V1^R^. Our model is of course not restricted to embryonic V1^R^, but may also apply to any electrically compact cell, the firing activity of which is dominated by *I*_*Nap*_ and delayed rectifier potassium currents. This is the case of many classes of embryonic cells in mammals at an early stage of their development. It can also apply to the axon initial segment, where *G*_*Nap*_ and *G*_*Kdr*_ are known to play the major role in the occurrence of repetitive firing [76]. Altogether our experimental and theoretical results provide a global view of the developmental trajectories of embryonic V1^R^ (see ***Figure 7F,G***). At E12.5, conductances of embryonic V1^R^ are widely spread in the *G*_*Nap*_− *G*_*Kdr*_ plane, which explains the heterogeneity of their firing patterns. This likely results from the random and uncorrelated expression of sodium and potassium channels from cell to cell at this early stage. Between E12.5 and E14.5-15.5 cell size increases, and *G*_*Kdr*_ with it, while the density of sodium channels decreases (see ***Figure 1* and *4***). The functional involution displayed by V1^R^ between E12.5 and E15.5 thus mainly results from a decrease of *G*_*Nap*_ coordinated with an increase of *G*_*Kdr*_. How these synergistic processes are controlled during this developmental period remains an open issue.

It is important to note that the presence of *I*_*Nap*_ is required for the functional diversity of V1^R^. Indeed, in the absence of *I*_*Nap*_, V1^R^ lose their ability to generate plateau potentials or to fire repetitively. More generally, when the diversity of voltage-gated channels is limited, as observed in embryonic neurons [18], changes in the balance between *I*_*Kdr*_ and non (or poorly) inactivating inward current can modify the firing pattern. This can be achieved not only by *I*_*Nap*_, but also by other slowly or non-inactivating inward conductances, such as *I*_*CAN*_ [75]. Our work also clearly indicates that a change in the firing pattern can only occur if a change in inward conductances cannot be counterbalanced by a corresponding change in outward conductances. This implies that there is no homeostatic regulation of channel density to ensure the robustness of V1^R^ excitability during its early development, contrarily to the mature CNS [37]. In addition, the poor repertoire of voltage-gated channels at this developmental stage precludes channel degeneracy, which is also known to ensure the robustness of excitability in mature neurons [37].

In conclusion, our results demonstrate that a simple mechanism involving only two slowly inactivating voltage-gated channels with opposite effects is sufficient to produce functional diversity in immature neurons having a limited repertoire of voltage-gated channels.

## Materials and Methods

### Isolated spinal cord preparation

Experiments were performed in accordance with European Community guiding principles on the care and use of animals (86/609/CEE, CE Off J no. L358, 18 December 1986), French decree no. 97/748 of October 19, 1987 (J Off République Française, 20 October 1987, pp. 12245-12248). All procedures were carried out in accordance with the local ethics committee of local Universities and recommendations from the CNRS. We used GAD67eGFP knock-in mice to visualize putative GABAergic INs [77], as in our previous study [25]. To obtain E12.5-E16.5 transgenic GAD67-eGFP embryos, 8 to 12 weeks old wild-type Swiss female mice were crossed with GAD67-eGFP Swiss male mice.

Isolated mouse SCs from 420 embryos were used in this work and obtained as previously described [28, 78]. Briefly, pregnant mice were anesthetized by intramuscular injection of a mix of ketamine and xylazine and sacrificed using a lethal dose of CO_2_ after embryos of either sex were removed. Whole SCs were isolated from eGFP-positive embryos and maintained in an artificial cerebrospinal fluid (ACSF) containing 135 mM NaCl, 25 mM NaHCO_3_, 1 mM NaH_2_PO_4_, 3 mM KCl, 11 mM glucose, 2 mM CaCl_2_, and 1 mM MgCl_2_ (307 mOsm/kg H_2_O), continuously bubbled with a 95% O_2_-5% CO_2_ gas mixture.

In the lumbar SC of GAD67eGFP mouse embryos, eGFP neurons were detected using 488 nm UV light. They were localized in the ventro-lateral marginal zone between the motor columns and the ventral funiculi [62]. Embryonic V1^R^ identity was confirmed by the expression of the forkhead transcription factor Foxd3 [25].

### Whole-cell recordings and analysis

The isolated SC was placed in a recording chamber and was continuously perfused (2 ml/min) at room temperature (22-26°C) with oxygenated ACSF. Whole-cell patch-clamp recordings of lumbar spinal embryonic V1^R^ were carried out under direct visualization using an infrared-sensitive CCD video camera. Whole-cell patch-clamp electrodes with a resistance of 4-7 MΩ were pulled from thick-wall borosilicate glass using a P-97 horizontal puller (Sutter Instrument Co., USA). They were filled with a solution containing (in mM): 96.4 K methanesulfonate, 33.6 KCl, 4 MgCl_2_, 4 Na_2_ ATP, 0.3 Na_3_ GTP, 10 EGTA, and 10 HEPES (pH 7.2; 290 mOsm/kg-H_2_O). This intracellular solution led to an equilibrium potential of chloride ions, *E*_*cl*_, of about -30 mV, close to the physiological values measured at E12.5 in spinal MNs [28]. The junction potential (6.6 mV) was systematically corrected offline.

Signals were recorded using Multiclamp 700B amplifiers (Molecular Devices, USA). Data were low-pass filtered (2 kHz), digitized (20 kHz) online using Digidata 1440A or 1550B interfaces and acquired using pCLAMP 10.5 software (Molecular Devices, USA). Analyses were performed off-line using pCLAMP 10.5 and Axograph 1.7.2 (Molecular devices; RRID:SCR_014284) software packages.

In voltage-clamp mode, voltage-dependent K^+^ currents (*I*_*Kv*_) were elicited in the presence of 1 μM TTX by 500 ms depolarizing voltage steps (10 mV increments, 10 s interval) after a prepulse of 300 ms at *V*_*H*_ = -100 mV. To isolate *I*_*Kdr*_, voltage steps were applied after a 300 ms prepulse at *V*_*H*_ = -30 mV that inactivated the low threshold transient potassium current *I_A_*. *I*_*A*_ was then obtained by subtracting offline *I*_*Kdr*_ from the total potassium current *I*_*Kv*_. Capacitance and leak current were subtracted using on-line P/4 protocol provided by pCLAMP 10.5.

In current-clamp mode, V1^R^ discharge was elicited using 2 s depolarizing current steps (from 0 to ≈ 50 pA in 5-10 pA increments, depending on the input resistance of the cell) with an 8 s interval to ensure that the membrane potential returned to *V*_*H*_. When a cell generated a sustained discharge, the intensity of the depolarizing pulse was reduced to the minimal value compatible with repetitive firing.

*I*_*Nap*_ was measured in voltage-clamp mode using a 70 mV/s depolarizing voltage ramp [79]. This speed was slow enough to preclude substantial contamination by the inactivating transient current and fast enough to avoid substantial inactivation of *I*_*Nap*_. Subtraction of the current evoked by the voltage ramp in the presence of 1 μM tetrodotoxin (TTX) from the control voltage ramp-evoked current revealed *I*_*Nap*_.

### Pharmacological reagents

During patch-clamp recordings, bath application of TTX (1 µM, Alomone, Israel) or 4-AP (0.3 - 600 µM, Sigma) was done using 0.5 mm diameter quartz tubing positioned, under direct visual control, 50 µm away from the recording area. The quartz tubing was connected to 6 solenoid valves linked with 6 reservoirs *via* a manifold. Solutions were gravity-fed into the quartz tubing. Their application was controlled using a VC-8 valve controller (Warner Instruments, USA).

4-aminopyridine (4-AP; Sigma Aldrich, USA) was used to block *I*_*Kdr*_. To determine the concentration–response curve, *I* − *V* curves of *I*_*Kdr*_ for different concentrations of 4-AP (0.3 to 300 μM) were compared to the control curve obtained in the absence of 4-AP. The percentage of inhibition for a given concentration was calculated by dividing the peak intensity of *I*_*Kdr*_ by the peak value obtained in control condition. The obtained normalized concentration–response curves were fitted using the Hill equation:

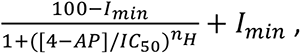

where [4-AP] is the 4-AP concentration, *I_min_* is the residual current (in percentage of the peak *I*_Kdr_), 100 − *I*_*min*_ is the maximal inhibition achieved for saturating concentration of 4- AP, *IC*_50_ is the 4-AP concentration producing half of the maximal inhibition, and *n*_*H*_ is the Hill coefficient. Curve fitting was performed using KaleidaGraph 4.5 (Synergy Software, USA).

### Immunohistochemistry and confocal microscopy

E14.5 embryos were collected from pregnant females. Once dissected out of their yolk sac, SCs were dissected and immediately immersion-fixed in phosphate buffer (PB 0.1 M) containing 4% paraformaldehyde (PFA; freshly prepared in PB, pH 7.4) for 1 h at 4°C. Whole SCs were then rinsed out in 0.12 M PB at 4°C, thawed at room temperature, washed in PBS, incubated in NH_4_Cl (50 mM), diluted in PBS for 20 min and then permeabilized for 30 min in a blocking solution (10% goat serum in PBS) with 0.2% Triton X-100. They were incubated for 48 h at 4°C in the presence of the following primary antibodies: guinea pig anti-FoxD3 (1:5000, gift from Carmen Birchmeier and Thomas Müller of the Max Delbrück Center for Molecular Medicine in Berlin) and rabbit anti-cleaved Caspase-3 (1:1000, Cell Signaling Technology Cat# 9661, RRID:AB_2341188). SCs were then washed in PBS and incubated for 2 h at RT with secondary fluorescent antibodies (goat anti-rabbit-conjugated 649; donkey anti-guinea pig-conjugated Alexa Fluor 405 [1:1000, ThermoFisher]) diluted in 0.2% Triton X-100 blocking solution. After washing in PBS, SCs were dried and mounted in Mowiol medium (Millipore, Molsheim, France). Preparations were then imaged using a Leica SP5 confocal microscope. Immunostaining was observed using a 40X oil-immersion objective with a numerical aperture of 1.25, as well as with a 63X oil-immersion objective with a numerical aperture of 1.32. Serial optical sections were obtained with a Z-step of 1 µm (40X) and 0.2-0.3 µm (63X). Images (1024x1024; 12-bit color scale) were stored using Leica software LAS-AF and analyzed using ImageJ 1.5 (N.I.H., USA; http://rsb.info.nih.gov/ij/) and Adobe Photoshop CS6 (Adobe, USA) software.

### Cluster analysis

To classify the firing patterns of embryonic V1^R^, we performed a hierarchical cluster analysis on a population of 163 cells. Each cell was characterized by three quantitative measures of its firing pattern (see legend of Figure 5). After normalizing these quantities to zero mean and unit variance, we performed a hierarchical cluster analysis using the hclust function in R software (version 3.3.2, https://cran.r-project.org/) that implements the complete linkage method. The intercluster distance was defined as the maximum Euclidean distance between the points of two clusters, and, at each step of the process, the two closest clusters were merged into a single one, thus constructing progressively a dendrogram. Clusters were then displayed in data space using the dendromat function in the R package ‘squash’ dedicated to color-based visualization of multivariate data. The best clustering was determined using the silhouette measure of clustering consistency [80]. The silhouette of a data point, based on the comparison of its distance to other points in the same cluster and to points in the closest cluster, ranges from -1 to 1. A value near 1 indicates that the point is well assigned to its cluster, a value near 0 indicates that it is close to the decision boundary between two neighboring clusters, and negative values may indicate incorrect assignment to the cluster. This allowed us to identify an optimal number k of clusters by maximizing the overall average silhouette over a range of possible values for k [80], using the silhouette function in the R package ‘cluster’.

### Biophysical modeling

To understand the relationship between the voltage-dependent membrane conductances and the firing patterns of embryonic V1^R^, we relied on a single compartment conductance-based model that included the leak current, the transient and persistent components of the sodium current, *I*_*Nat*_ and *I*_*Nap*_, a delayed rectifier potassium current *I*_*Kdr*_ and the inactivating potassium current *I*_*A*_ revealed by experiments. Voltage evolution then followed the equation

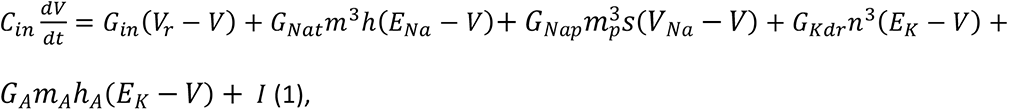

where *C*_*in*_ was the input capacitance; *G*_*in*_ the input conductance; *G*_*Nat*_, *G*_*Nap*_, *G*_*Kdr*_ and *G*_*A*_ the maximal conductances of the aforementioned currents; *m*, *m*_*p*_, *n* and *m*_*A*_ their activation variables; ℎ the inactivation variable of *I*_*Kdr*_, *s* the slow inactivation variable of *I*_*Nat*_, and *m*_*A*_ the inactivation variable of *I*_*A*_. *V*_*r*_ is the baseline potential imposed by *ad hoc* current injection in current-clamp experiments; *E*_*Na*_ and *E*_*K*_ are the Nernst potentials of sodium and potassium ions, and *I* the injected current. All gating variables satisfied equations of the form:

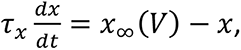

where the (in)activation curves were modeled by a sigmoid function of the form:

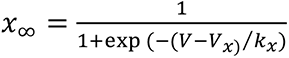

with *k_x_*being positive for activation and negative for inactivation. The time constant τ_*s*_ was voltage-independent except for the inactivation variables ℎ and *s*. The activation variable *m*_*A*_ of *I*_*A*_ was assumed to follow instantaneously voltage changes.

The effect of channel noise was investigated with a stochastic realization of the model, where channels kinetics were described by Markov-type models, assuming a unitary channel conductance of 10 pS for all channels.

### Choice of model parameters

Most model parameters were chosen on the basis of experimental measurements performed in the present study or already reported [25]. Parameters that could not be constrained from our experimental data were chosen from an experimentally realistic range of values. *V*_*r*_ was set at -60 mV as in experiments (see ***table 1***). *C*_*in*_ (average 13.15 pF, 50% between 11.9 and 15.1 pF, only 18 cells out of 246 in the first quartile below 7.2 pF or in the fourth quartile above 19 pF) and *G*_*in*_ (50% of cells between 0.71 and 1.18 nS, only 7 out of 242 with input conductance above 2 nS) were not spread much in the cells recorded at E12.5, which showed that most embryonic V1^R^ were of comparable size. Interestingly, *C*_*in*_ and *C*_*in*_ were not correlated, which indicated that the input conductance was determined by the density of leak channels rather than by the sheer size of the cell. Moreover, no correlation was observed between the passive properties and the firing pattern [25]. Therefore, we always set *C*_*in*_ and *C*_*in*_ to 1 nS and 13 pF in the model (except in ***Figure 6––figure supplement 2***), close to the experimental medians (0.96 nS and 13.15 pF, respectively). The membrane time constant *C*_*in*_/*G*_*in*_ was then equal to 13 ms, which was also close to the experimental median (13.9 ms, N=241). *E*_*Na*_ was set to 60 mV (see [25]). The activation curve of *I*_*Kdr*_ was obtained by fitting experimental data, leading to an average mid-activation of -36 mV and an average steepness of 9.5 mV. The experimentally measured values of *G*_*Kdr*_ were in the range 0-2.2 nS. We assumed that the activation curve of *I*_*Kdr*_ was shifted rightward by 10 mV in comparison to *I*_*Kdr*_. No experimental data was available for the inactivation of *I*_*Kdr*_. We chose a mid-inactivation voltage *V*_*H*_= -45 mV and a steepness *k_h_* =-5 mV. We also assumed that the activation time constant of both *I*_*Nat*_ and *I*_*Nav*_ was 1.5 ms, and that the inactivation time constant was voltage-dependent: τ_*s*_*V* = 16.5 − 13.5 tanh((*V* + 20) 15), decreasing by one order of magnitude (from 30 ms down to 3 ms) with the voltage increase. This enabled us to account for the shape of the action potentials recorded in experiments, showing a slow rise time and rather long duration. The conductance *G*_*Kdr*_ was not measured experimentally. When choosing a reasonable value of 20 nS for *G*_*Nat*_, the model behaved very much as recorded embryonic V1^R^: with similar current threshold (typically 10-20 pA) and stable plateau potential obtained for the largest values of *G*_*Nap*_.

**Table 1.**

| Parameter | Basic model | Model with slow inactivation<br>of $I_{Nap}$ |
| --- | --- | --- |
| Passive parameters |  |  |
| Input conductance $G_{in}$ | 1 nS | same |
| Input capacitance $C_{in}$ | 13 pF (E12.5, Figs. 7B, C, D and F and 8B to D) or 18 pF (E14.5, Fig. 7G) | 13 pF |
| Resting potential $V_r$ | -60 mV | same |
| Injected current $I$ | 20 pA (Fig. 7B to G) | 10 pA (Fig. 8C) or 12 pA (Fig. 8D)<br>variable in Fig. 8B |
| Transient sodium current $I_{nat}$ | | |
| Maximal conductance $G_{Nat}$ | 20 nS | same |
| Reversal potential $E_{Na}$ | 60 mV | |
| Activation exponent | 3 |  |
| Mid-activation $V_m$ | -26 mV | |
| Steepness of activation $k_m$ | 9.5 mV | |
| Activation time constant | 1.5 ms |  |
| Mid-inactivation $V_h$ | -45 mV | |
| Steepness of inactivation $K_h$ | -5 mV | |
| Inactivation time constant $\tau_m$ | Voltage-dependent (see Material and Methods) | |
| Persistent sodium current $I_{Nap}$ | | |
| Maximal conductance | variable (see text and figure captions) | same |
| Mid-activation voltage | -36 mV | same |
| Mid-inactivation $V_s$ | | -30 mV |
| Steepness of inactivation $k_s$ | | -5 mV |
| Inactivation time constant | Slow inactivation not included | 2 s |
| Delayed rectifier potassium current $I_{Kdr}$ | | |
| Maximal conductance $G_{Kdr}$ | variable (see text and figure captions) | same |
| Reversal potential $E_K$ | -96 mV | |
| Activation exponent | 3 |  |
| Mid-activation $V_n$ | -20 mV | |
| Steepness of activation $k_n$ | 15 mV | |
| Activation time constant $\tau_m$ | 10 ms | |
| Potassium A current $I_A$ (when included in the basic model) | | |
| Maximal conductance $G_A$ | Equal to $G_{Kdr}$ | never included |
| Mid-activation $V_{mA}$ | -30 mV | |
| Steepness of activation $k_{mA}$ | 12 mV | |
| Activation time constant | Instantaneous activation |  |
| Mid-inactivation $V_{hA}$ | -70 mV | |
| Steepness of inactivation $k_{hA}$ | -7 mV | |
| Inactivation time constant $\tau_{hA}$ | 23 ms | |

When taking into account slow inactivation of *I*_*Nap*_ (see ***Figure 8***), we chose *V*_*s*_ = -30 mV for the mid-inactivation voltage and set the steepness *k_s_* at -5 mV (as for the inactivation of *I*_*Nat*_). For simplicity, we assumed that the inactivation time constant was voltage-independent and set it at a value of 2 s.

*E*_*K*_ was set to the experimentally measured value of -96 mV [25]. The activation parameters of *I*_*Kdr*_ were obtained by fitting the experimental data: *V*_*n*_ = -20 mV, *k_n_* = 15 mV, τ_*n*_= 10 ms and an activation exponent of 3. The activation and inactivation properties of *I*_*A*_ were also chosen based on experimental measurements. Accordingly, *V*_*m_A_*_= -30 mV, *k*_*m_A_*_= -12 mV, *V*_*h*_*A*__= -70 mV, *k*_*h*_*A*__ = -7 mV, and τ_*h*_*A*__= 23 ms. When *I*_*A*_ was taken into account, we assumed that *G*_*A*_= *G*_*Kdr*_ consistently with experimental data (see ***Figure 6––figure supplement 1***).

### Numerical simulations and dynamical systems analysis

We integrated numerically the deterministic model using the freeware XPPAUT [81] and a standard fourth-order Runge-Kutta algorithm. XPPAUT was also used to compute one-parameter and two-parameters bifurcation diagrams. The stochastic version of the model was also implemented in XPPAUT and computed with a Gillespie’s algorithm [82].

To investigate the dynamics of the model with slow inactivation of *I*_*Nav*_, we relied on numerical simulations together with fast/slow dynamics analysis [83]. In this approach, one distinguishes slow dynamical variables (here only *s*) and fast dynamical variables. Slow variables vary little at the time scale of fast variables and may therefore be considered as constant parameters of the fast dynamics in first approximation. In contrast, slow variables are essentially sensitive to the time average of the fast variables, much more than to their instantaneous values. This separation of time scales allows one to conduct a phase plane analysis of the full dynamics.

### Statistics

Samples sizes (n) were determined based on previous experience. The number of embryos (N) is indicated in the main text and figure captions. No power analysis was employed, but sample sizes are comparable to those typically used in the field. All values were expressed as mean with standard error of mean (SEM). Statistical significance was assessed by non-parametric Kruskal-Wallis test with Dunn’s post hoc test for multiple comparisons, Mann-Whitney test for unpaired data or Wilcoxon matched pairs test for paired data using GraphPad Prism 7.0e Software (USA). Significant changes in the proportions of firing patterns with age were assessed by chi-square test for large sample and by Fisher’s exact test for small sample using GraphPad Prism 7.0e Software. Significance was determined as p<0.05 (*), p<0.01 (**) or p<0.001 (***). The exact p value was mentioned in the result section or in the figure captions.

## Acknowledgments

We thank Susanne Bolte, Jean-François Gilles and France Lam for assistance with confocal imaging (IBPS imaging facility) and IBPS rodent facility team for animal care and production. We thank University Paris Descartes for hosting Yulia Timofeeva as an invited professor. This work was supported by INSERM, CNRS, Sorbonne Université (Paris), Université de Bordeaux, Université Paris Descartes and Fondation pour la Recherche Médicale.

## Supplementary legends

**Figure 2––figure supplement 1.**
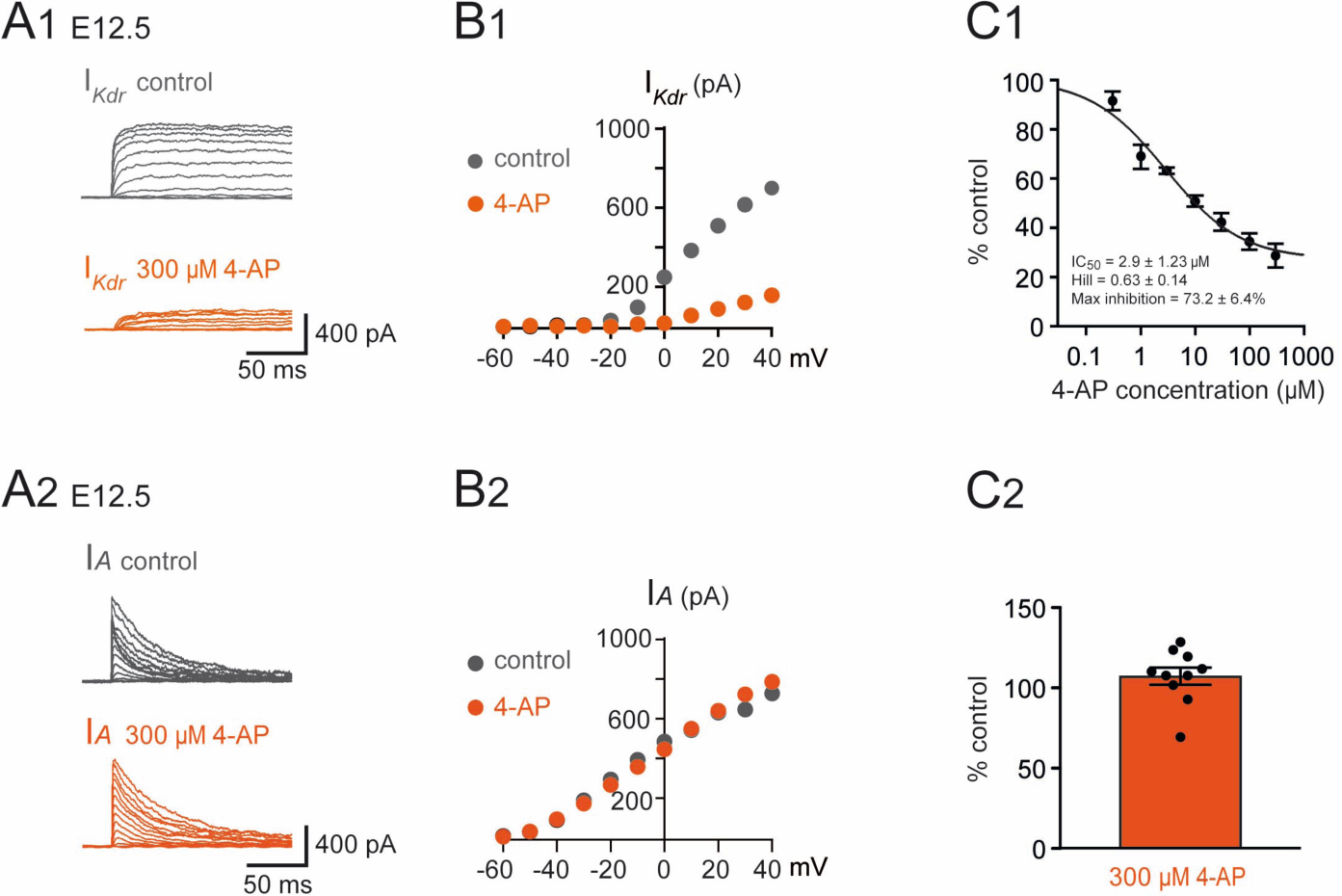
Effect of 4-AP on *I*_*Kdr*_ and *I_A_* in embryonic V1^R^. (A1) Example of voltage-dependent potassium currents evoked in response to 10 mV membrane potential steps (200 ms) from -100 mV or from -30 mV to +40 mV (10 s interval between pulses). V1 ^R^ were voltage clamped at *V*_*H*_ = -60 mV. A prepulse of -40 mV (300 ms) was applied to activate both *I*_*A*_ and *I*_*Kdr*_. *I*_*Kdr*_ was evoked in response to 10 mV membrane potential steps (200 ms) from -100 mV to +40 mV. V1 ^R^ were voltage clamped at *V*_*H*_ = -60 mV. A prepulse of 30 mV (*V*_*H*_ = - 30 mV) was applied to isolate *I*_*Kdr*_. (A1) Representative example of the effect of 300 μM 4-AP application on *I*_*Kdr*_ recorded from embryonic V1^R^ at E12.5. (B1) Curves showing current-voltage relationships of *I*_*Kdr*_ in control and in the presence of 300 μM 4-AP. Measurements were performed on traces shown in A1. (C1) Dose-response relationship of 4-AP-evoked *I*_*Kdr*_ inhibition (mean + SE). Data were normalized to *I*_*Kdr*_ amplitude measured in the absence of 4-AP (*V*_*H*_ = 40mV) and fitted as explained in Materials and Methods. Note that 4-AP IC_50_ is in μM range (2.9 μM). 0.3 μM 4-AP n = 3; N = 3, 1 μM 4- AP n = 3; N = 3, 3 μM 4-AP n = 9; N = 9, 10 μM 4-AP n = 13; N = 13, 30 μM 4-AP n = 7; N = 7, 100 μM 4-AP n = 7; N = 7, 300 μM 4-AP n = 7; N = 7. (A2) *I_A_* was obtained as the difference between currents evoked from *V*_*H*_ = -100 mV and currents evoked from *V*_*H*_ = -30 mV (10 mV voltage step). (A2) Representative example of the effect of 300 μM 4-AP on *I*_*A*_ in V1^R^ recorded at E12.5. (B2) *I*_*A*_ Current-voltage (*I* − *V*) relationship in control conditions and in the presence of 300 μM 4-AP. The *I* − *V* curves were obtained from the traces shown in A1. (C2) Bar graph showing the percentage of *I*_*A*_ block elicited by 4-AP. Note that 4-AP did not significantly block *I*_*A*_(Wilcoxon test *P* = 0.065, n = 10).

**Figure 2––figure supplement 2.**
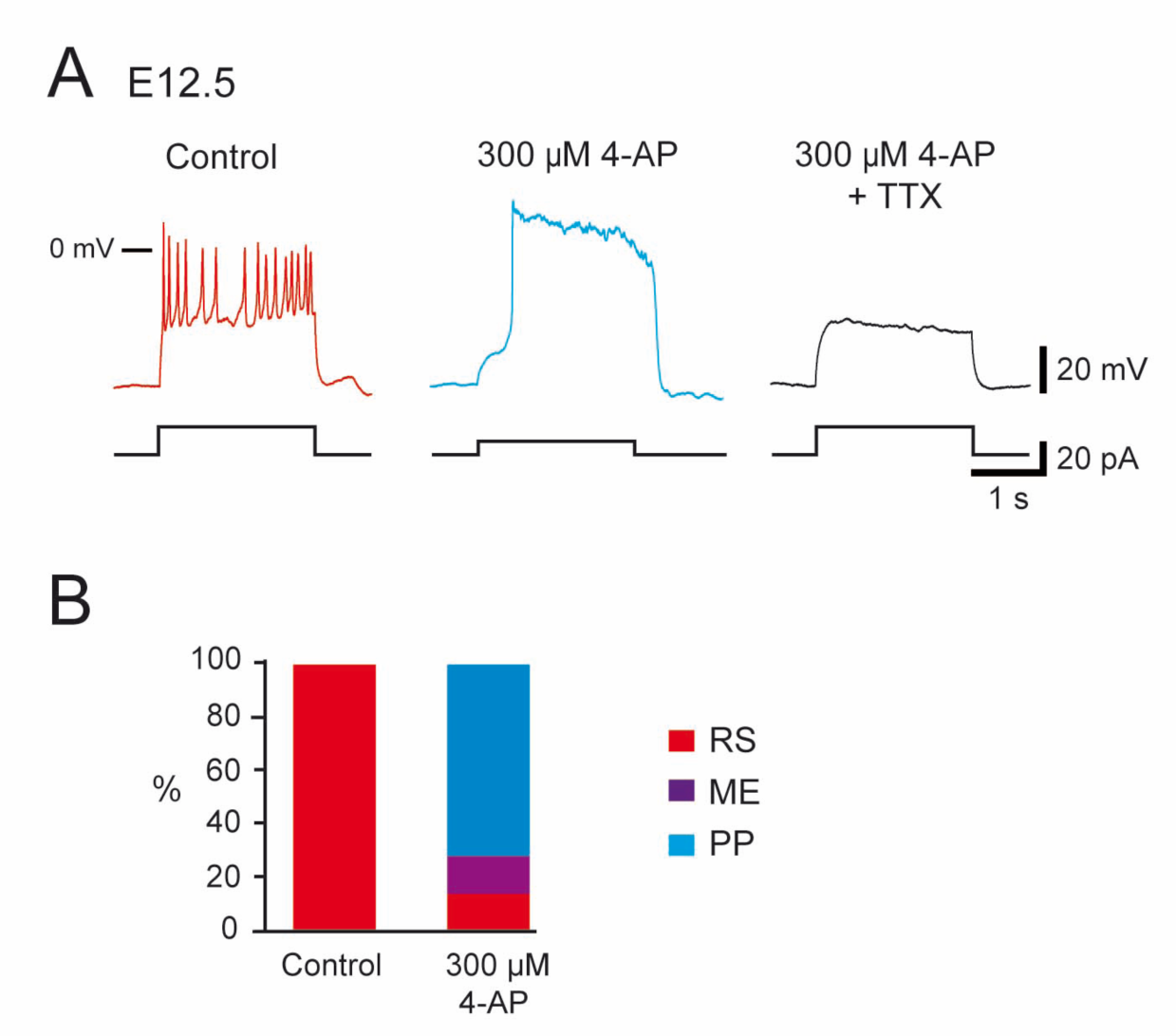
Relates to Fig 2. Effect of 4-AP application in repetitively spiking V1^R^ at E12.5. (A) Representative traces showing the effect of 4-AP application (300 μM) on Repetitive Spiking (RS) V1^R^ at E12.5. Note that plateau potential activity evoked in the presence of 4-AP (middle trace) was blocked by 0.5 μM TTX (right trace). (B) Bar plots showing the changes in the firing pattern of RS V1^R^ evoked by 300 μM 4-AP application (n = 14). 4-AP application evoked a plateau potential in 71.4 % of the recorded neurons (10/14) and mixed events in 14.3% of the recorded neurons (2/14). The excitability pattern was not modified in 2 neurons. Repetitive Spiking (RS) V1^R^ (red), Mixed events (ME) V1^R^ (purple), Plateau Potential (PP) V1^R^ (blue).

**Figure 3––figure supplement 1.**
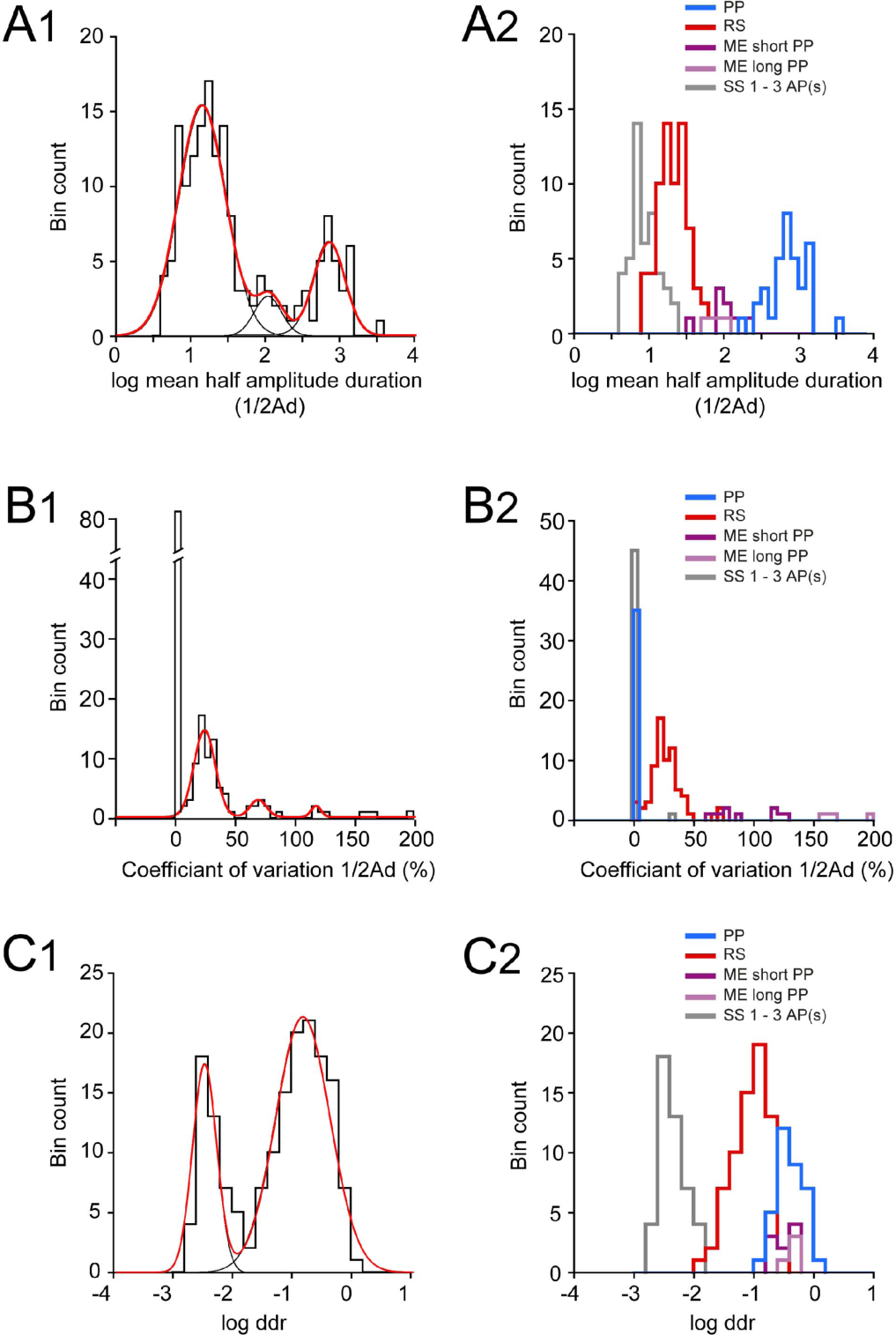
Distributions of log ½Ad, CV ½Ad and log ddr values related to the cluster analysis of embryonic V1^R^ firing patterns. (A1) Histogram of log mean ½Ad (mean half amplitude event duration) for the whole V1^R^ population at E12.5 (n= 164; bin width 0.1). The histogram was well fitted by the sum of three Gaussian curves with means and SDs of 1.135, 2.046 & 2.84, and 0.316, 0.181 & 0.21, respectively. (A2) Histogram of the values of log mean ½Ad sorted after cluster analysis showing single spiking (SS) V1^R^ (gray), repetitive spiking (RS) V1^R^ (red), mixed events (ME) V1^R^ with short plateau potentials (ME short PP V1^R^, light purple), ME V1^R^ with long plateau potentials (ME long PP V1^R^, dark purple) and plateau potential (PP) V1^R^ (blue). log mean ½Ad was significantly different between SS V1^R^, PP V1^R^, the whole ME V1^R^ population (ME_s_ and ME_l_ V1^R^) and PP V1^R^ (Kruskall-Wallis test *P* < 0.0001; SS V1^R^ versus RS V1^R^, *P* < 0.0001; SS V1^R^ versus ME V1^R^, *P* < 0.0001; SS V1^R^ versus PP V1^R^, *P* < 0.0001; RS V1^R^ versus ME V1^R^, *P* = 0.0004; RS V1^R^ versus PP V1^R^, *P* < 0.0001; ME V1^R^ versus PP V1^R^, *P* = 0.018; SS V1^R^ n = 46, RS V1^R^ n = 69, ME_s_ V1^R^ n = 9, ME_l_ V1^R^ n = 4, PP V1^R^ n = 35). (B1) Histogram of CV ½Ad for the whole V1^R^ population at E12.5 (n= 164; bin width 5%). Note that a large population of V1^R^ had zero CV ½Ad (n = 83). The histogram for CV ½Ad ≠ 0 was fitted by the sum of three Gaussian curves with means and SDs of 23.4, 68.4 & 117 (%) and 8.9, 6.8 & 4.1, respectively. (B2) Histograms of the values of CV ½Ad sorted after cluster analysis showing SS V1^R^ (black), RS V1^R^ (red), ME_s_ V1^R^ (light purple), ME_l_ V1^R^ (dark purple) and PP V1^R^. CV ½Ad was not significantly different between SS V1^R^ and PP V1^R^ (CV ½Ad of SS V1^R^ and PP V1^R^ = 0.682 % and 0% respectively: only one of the 46 SS V1^R^ displayed 3 PA and had a CV ½Ad of 31.37). CV ½Ad was significantly different between RS V1^R^ and the whole ME V1^R^ population and also between SS V1^R^ or PP V1^R^ and RS V1^R^ or ME V1^R^ (Kruskall-Wallis test *P* < 0.0001; SS V1^R^ versus RS V1^R^ *P* < 0.0001, SS V1^R^ versus ME V1^R^ *P* < 0.0001, SS V1^R^ versus PP V1^R^ *P* = 0.846, RS V1^R^ versus ME V1^R^ *P* = 0.0003, RS V1^R^ versus PP V1^R^ *P* < 0.0001, ME V1^R^ versus PP V1^R^ *P* < 0.0001). (C1) Histogram of log ddr (sum of ½Ad divided by pulse duration) for the whole V1^R^ population at E12.5 (n= 164; bin width 0.2). The histogram was fitted by the sum of two Gaussian curves with means and SDs of -2.51 & -0.851, and 0.2 & 0.46, respectively. (C2) Histograms of the values of log ddr sorted after cluster analysis showing SS V1^R^ (black), RS V1^R^ (red), ME_s_ V1^R^ (light purple), ME_l_ V1^R^ (dark purple) and PP V1^R^. log (ddr) was not significantly different between ME V1^R^ and PP V1^R^, while it was significantly different between SS V1^R^ and RS V1^R^, SS V1^R^ and the whole ME V1^R^ population, SS V1^R^ and PP V1^R^, RS V1^R^ and the whole ME V1^R^ population, RS V1^R^ and PP V1^R^ (Kruskall-Wallis test *P* < 0.0001; SS V1^R^ versus RS V1^R^, *P* < 0.0001; SS V1^R^ versus ME V1^R^, *P* < 0.0001; SS V1^R^ versus PP V1^R^, *P* < 0.0001; RS V1^R^ versus ME V1^R^, *P* < 0.0001; RS V1^R^ versus PP V1^R^, *P* < 0.0001; ME V1^R^ versus PP V1^R^, *P* = 0.977). ME_s_ V1^R^ and ME_l_ V1^R^ differed only by their CV ½Ad (Mann-Whitney test, log mean ½Ad for ME_s_ V1^R^ versus log mean ½Ad for ME_l_ V1^R^, *P* = 0.26; CV ½Ad for ME_s_ V1^R^ versus CV ½Ad ME_l_ V1^R^, *P* = 0.0028 and log ddr for ME_s_ V1^R^ versus log ddr for ME_l_ V1^R^, *P* = 0.1483). It is noteworthy that the distribution of the values of each metric was multimodal thus indicating that each of them could partially discriminate different groups of embryonic V1^R^ according to their firing pattern.

**Figure 6––figure supplement 1.**
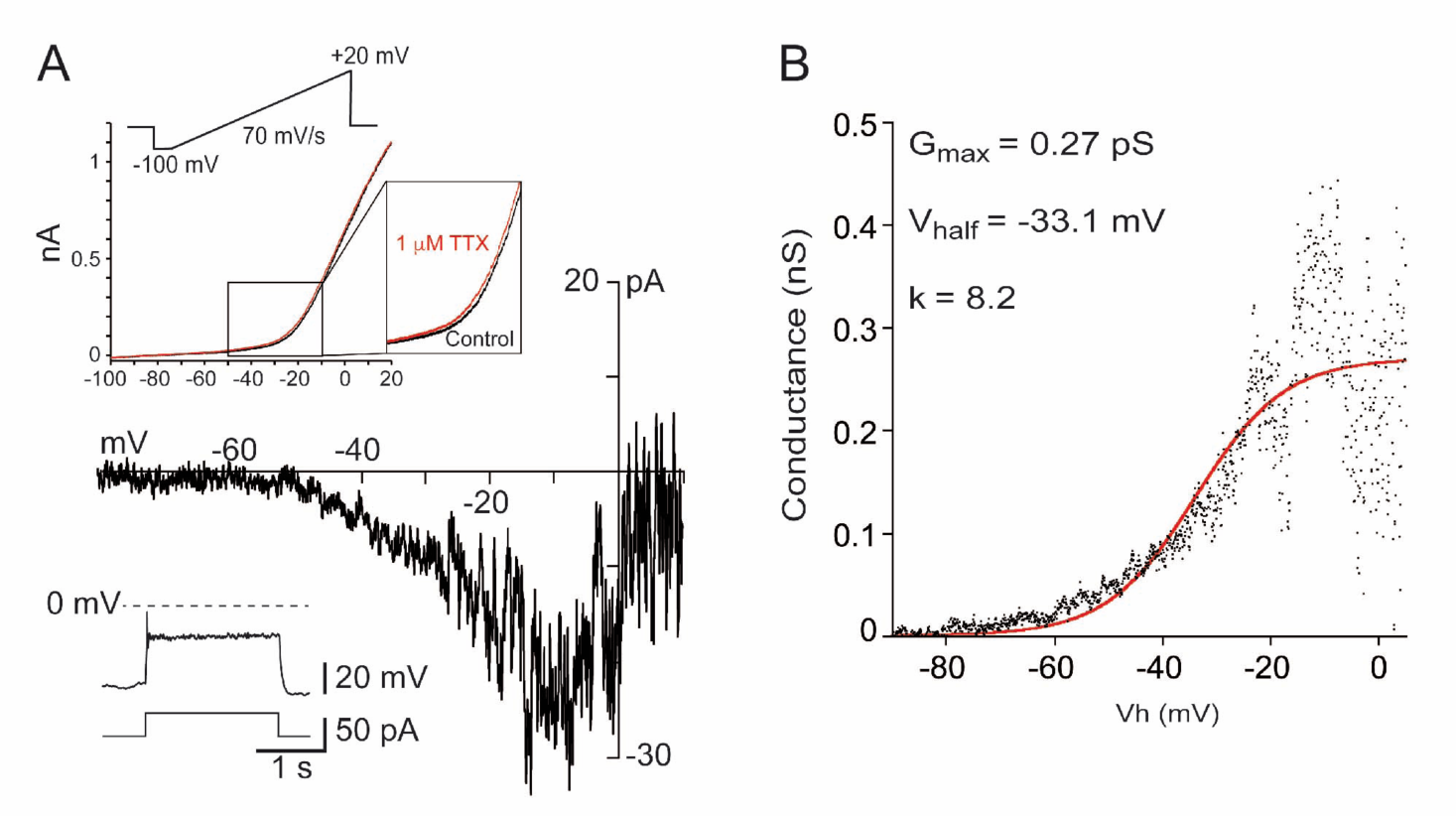
*I*_*Nap*_ is present in embryonic V1^R^ recorded at E14.5. (A) Representative trace of *I*_*Nap*_ evoked by a slow depolarizing voltage ramp (70 mV/s, upper insert) in SS embryonic V1^R^ (lower insert). *I*_*Nap*_ was isolated by subtracting currents evoked by depolarizing ramps in the presence of 1 μM TTX to the control current evoked in the absence of TTX (upper insert). (B) Voltage dependence of G*_Nap_* conductance calculated from the trace shown in A. The activation curve was obtained by transforming the current evoked by a depolarizing voltage ramp from -100 mV to 20 mV (70 mV/s) using the following equation: G_NaP_ = -I_Nap_/((-Vh)+E_Na_) where Vh is the holding potential at time t during a depolarizing voltage ramp and E_Na_ is the equilibrium potential for sodium (E_Na_ = 60 mV). The G_NaP_/Vh curve was fitted with the following Boltzmann function: G = G_MAX_/(1+exp(-(V- V_HALF_)/k))) (Boeri et al. 2018), where V_half_ is the Vh value for G_Nap_ half activation, k the slope factor of the curve and G_max_ the maximum conductance. We found no significant difference between the values of V_half_ (Mann-Whitney test: *P* = 0.8518) and of k (Mann-Whitney test: *P* = 0.7546) obtained at E12.5 (Boeri et al. 2018) and those obtained at E14.5. At E14.5 V_half_ = - 27 ± 5.1 mV and k = 7.73 ± 0.78 (n = 6).

**Figure 6––figure supplement 2.**
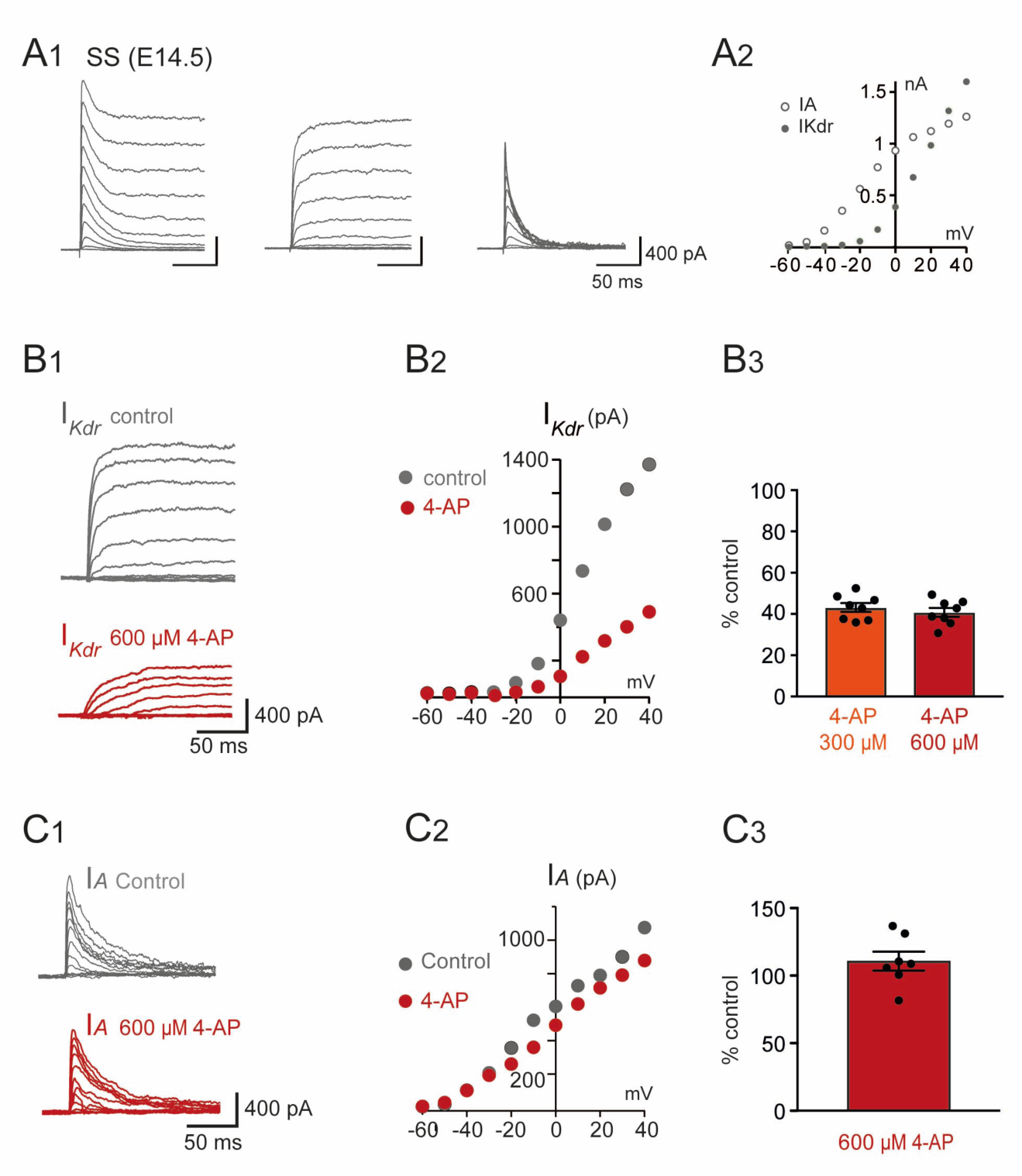
*I*_*Kdr*_ was inhibited by 4-AP in V1^R^ recorded at E14.5. (A1) Representative examples of the total outward K^+^ currents obtained from *V*_*H*_ = -100 mV (left traces), of *I*_*Kdr*_ (*V*_*H*_ = -30 mV, middle traces) and of isolated *I*_*A*_(left traces) recorded in single spiking (SS) V1^R^ at E14.5. (A2) Current-voltage relationship of *I*_*Kdr*_ (filled circle) and of *I*_*A*_(open circle) in SS V1^R^ at E14.5. *I* − *V* curves were obtained from currents shown in A1. (B1) Representative example of the effect of 4-AP at 600 μM in V1^R^ at E14.5. (B2) Current- voltage curves in control condition and in the presence of 600 μM 4-AP. (B3) Bar plots showing the percentage of *I*_*Kdr*_ inhibition evoked by 300 μM 4-AP application (n = 8) and by 600 μM 4-AP application (n = 7). The percentages of *I*_*Kdr*_ inhibition evoked by 300 μM 4-AP and by 600 μM 4-AP applications were not significantly different (*P* = 0.574). (C1) Representative example of the effect of 600 μM 4-AP on *I*_*A*_ in V1^R^ recorded at E14.5. (C2) *I* − *V* curves in control conditions and in the presence of 600 μM 4-AP. These curves were obtained from the traces shown in B1. (C3) Bar graph showing the percentage of *I*_*A*_ block elicited by 4-AP. 4-AP did not significantly block *I*_*A*_(Wilcoxon test *P* = 0.11, n = 6).

**Figure 7––figure supplement 1.**
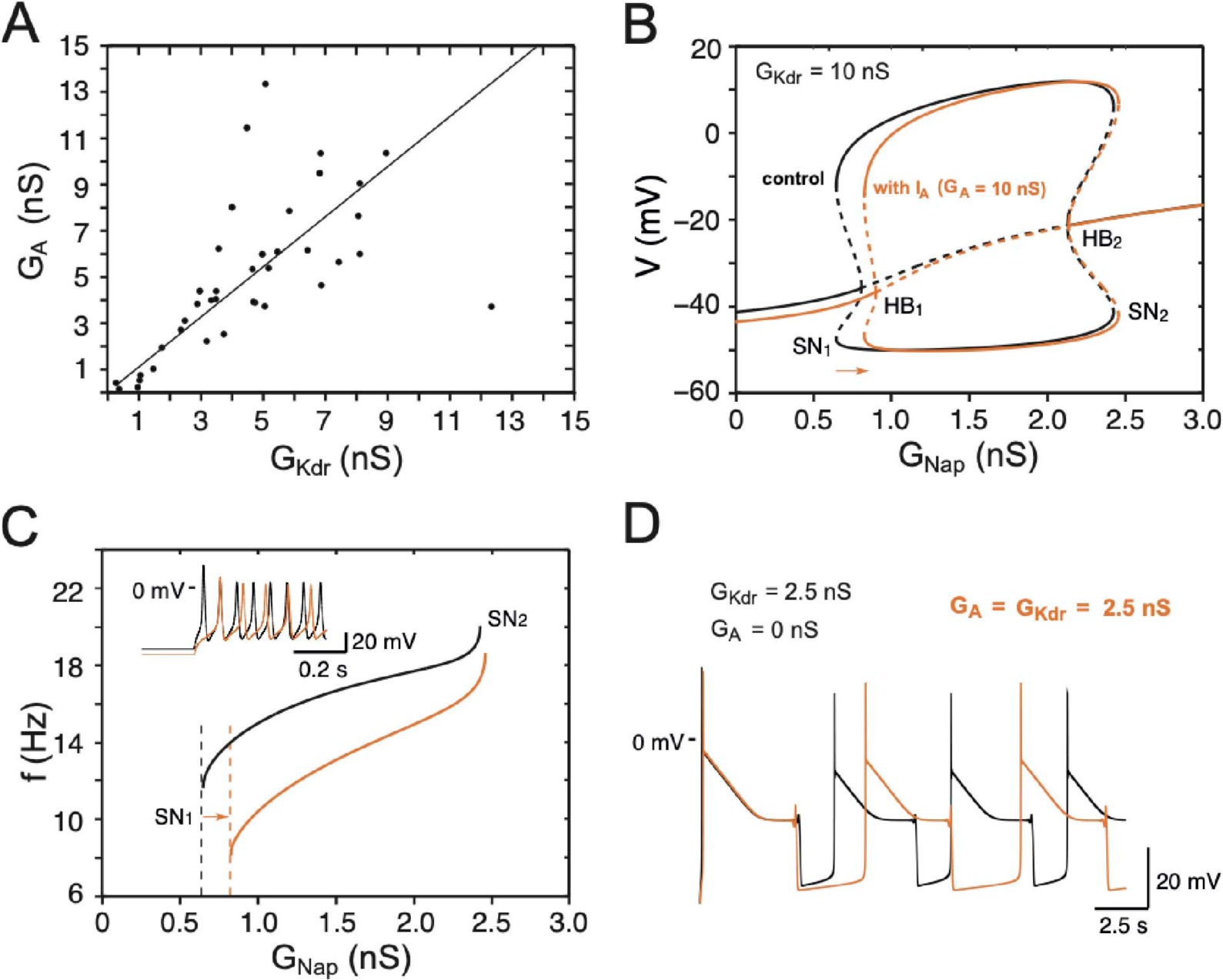
Effects of *I*_*A*_ on embryonic V1^R^ firing patterns predicted by computational modeling. (A) The maximal conductances of *I*_*Kdr*_ and *I*_*A*_ at E12.5 are linearly correlated. Best fit: *G*_*A*_= 1.09 *G*_*Kdr*_(R^2^ = 0.81, N=44). (B) Effect of *I*_*A*_ on the dynamics of the basic model. The one- parameter bifurcation diagrams in control condition (black, *I* = 20 pA, *G*_*Kdr*_= 10 nS, no *I*_*A*_, same as in Fig 7B) and with *I*_*A*_ added (orange, *G*_*A*_= 10 nS) are superimposed. The *I*_*A*_ current shifts the firing threshold SN_1_ to the right by 0.18 nS (see also C) as indicated by the orange arrow, with little effect on the amplitude of action potentials (see also insert in C). In contrast, *I*_*A*_ shifts SN_2_ by only 0.03 nS because it is inactivated by depolarization. (C) *I*_*A*_ also slows down the discharge frequency, as shown by comparing the *G*_*Nap*_− *V* curves without *I*_*A*_(black) and with *I*_*A*_(orange). For *G*_*Nap*_= 1 nS, for instance, the firing frequency is reduced by 31%, from 15 to 10.4 Hz. Here again, the effect of *I*_*A*_ progressively decreases as *G*_*Nap*_ increases because of the membrane depolarization elicited by *I*_*Nap*_. For *G*_*Nap*_= 2.4 nS, for instance, the firing frequency is reduced by 11% only, from 19.1 to 17 Hz. This frequency reduction elicited by *I*_*A*_ does not merely result from the increased firing threshold. Note also that the latency of the first spike is increased (see voltage trace in insert), which is a classical effect of *I*_*A*_. (D) *I*_*A*_ reduces the frequency of pseudo-plateau bursting by lengthening quiescent episodes (doubling their duration in the example shown) without affecting the duration of plateaus much (here a mere 5% increase), as shown by the comparison of the voltage traces obtained without *I*_*A*_(control, *G*_*Kdr*_= *2*.5 nS, black) and with *I*_*A*_(*G*_*Kdr*_= *G*_*A*_= *2.5* nS, orange). This is because *I*_*A*_ is activated near rest but inactivated during voltage plateaus. Note that increasing *G*_*Kdr*_, in the absence of *I*_*A*_ has not the same effect; it shortens both plateaus and quiescent episodes (see Fig 8C, where *G*_*Kdr*_= 5 nS). Again, this is because *I*_*Kdr*_ does not inactivate (or does it only very slowly), in contrast to *I*_*A*_.

**Figure 7––figure supplement 2.**
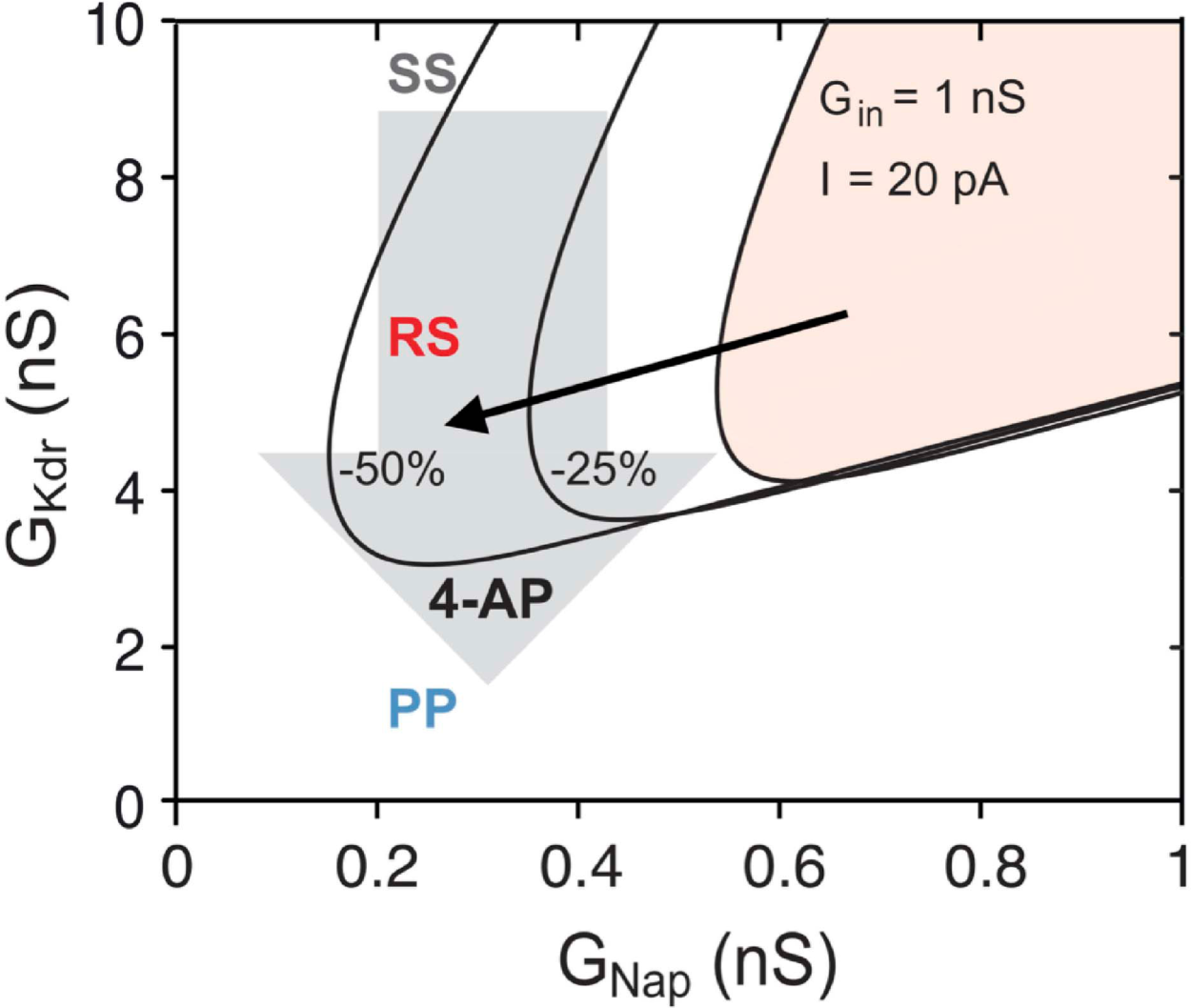
Explaining the effect of 4-AP on the firing pattern. The RS region of the basic model, where repetitive firing may occur, is displayed in the *G*_*Nap*_ – *G*_*Kdr*_ plane in control condition for E12.5 V1^R^ (*C*_*in*_= 13 pF, *G*_*in*_= 1 nS, *I* = 20 pA, shaded area), and when *C*_*in*_ and *I* were both reduced by 25% (middle curve) or by 50% (left curve). The reduced *I* accounts for the decrease in rheobase, and thus in the current injected in the experiments, following the decrease in *G*_*in*_. If 4-AP reduced only *G*_*Kdr*_(as indicated by the downward arrow) the firing pattern of SS V1^R^ would not change, the RS region being too far to the right to be visited. In contrast, when the effects of 4-AP on the input conductance and rheobase are taken into account, the bifurcation diagram moves leftwards and downwards, as indicated by the oblique black arrow, and the RS and PP regions are then successively entered as *G*_*Kdr*_ is reduced. The same explanation holds at E14.5.

